# Single-cell profiling and zebrafish avatars reveal *LGALS1* as immunomodulating target in glioblastoma

**DOI:** 10.1101/2023.04.27.538517

**Authors:** Lise Finotto, Basiel Cole, Wolfgang Giese, Elisabeth Baumann, Annelies Claeys, Maxime Vanmechelen, Brecht Decraene, Marleen Derweduwe, Nikolina Dubroja Lakic, Gautam Shankar, Madhu Nagathihalli Kantharaju, Jan Philipp Albrecht, Ilse Geudens, Fabio Stanchi, Keith L. Ligon, Bram Boeckx, Diether Lambrechts, Kyle Harrington, Ludo Van Den Bosch, Steven De Vleeschouwer, Frederik De Smet, Holger Gerhardt

## Abstract

Glioblastoma (GBM) remains the most malignant primary brain tumor, with a median survival rarely exceeding 2 years. Tumor heterogeneity and an immunosuppressive microenvironment are key factors contributing to the poor response rates of current therapeutic approaches. GBM-associated macrophages (GAMs) often exhibit immunosuppressive features that promote tumor progression. However, their dynamic interactions with GBM tumor cells remain poorly understood. Here, we used patient-derived GBM stem cell cultures and combined single-cell RNA sequencing of GAM-GBM co-cultures and real-time *in vivo* monitoring of GAM-GBM interactions in orthotopic zebrafish xenograft models to provide insight into the cellular, molecular, and spatial heterogeneity. Our analyses revealed substantial heterogeneity across GBM patients in GBM-induced GAM polarization and the ability to attract and activate GAMs – features that correlated with patient survival. Differential gene expression analysis, immunohistochemistry on original tumor samples, and knock-out experiments in zebrafish subsequently identified *LGALS1* as a primary regulator of immunosuppression. Overall, our work highlights that GAM-GBM interactions can be studied in a clinically relevant way using co-cultures and avatar models, while offering new opportunities to identify promising immune-modulating targets.

## Introduction

Glioblastoma (GBM) is the most common and aggressive type of malignant brain tumors in adults (Grochans *et al*, 2022). It is associated with a poor prognosis: a median survival of 15 months under standard of care (SoC) treatment, which comprises maximal safe surgical resection, followed by radiotherapy and chemotherapy (Stupp *et al*, 2005; Grochans *et al*, 2022). The main challenges presented by this malignancy are the cellular and molecular heterogeneity, the aggressive nature and invasive behavior, the inability to surgically resect the entire tumor, and the limitations of drug administration (Harder *et al*, 2018). Together, these problems invariably result in tumor progression or recurrence, demonstrating the urgent need for more effective treatments (Shergalis *et al*, 2018).

Cellular heterogeneity remains an important hallmark of GBM: it not only afflicts tumor cells, but is also present in cells of the tumor microenvironment (TME). GBM-associated macrophages (GAMs) represent 30-50% of the tumor and consist of tissue-resident microglia and tumor-infiltrating macrophages. Nevertheless, it remains unclear how GAM behavior is affected by tumor cell heterogeneity and how this contributes to GBM progression (Reimunde *et al*, 2021). In general, immunosuppressive tumor-associated macrophages (TAMs) are critical regulators of tumor progression, metastasis and immune evasion, and therefore promising therapeutic targets (Buonfiglioli & Hambardzumyan, 2021; Wang *et al*, 2022). In the last decade, therapeutic targeting of immune cells has gained great interest and impressive results have been achieved with immunotherapy in various types of cancer (Esfahani *et al*, 2020). However, GBM is a “cold” tumor and the limited success rate of immunotherapy relates to the highly immunosuppressive nature of the GBM microenvironment which is largely driven by GAMs and results in low levels of tumor-infiltrating lymphocytes and T cell exhaustion (Yu & Quail, 2021). Despite significant scientific efforts, the SoC treatment for GBM has not changed for more than 15 years. Therefore, landscaping the levels and types of GAM polarization across the GBM patient population could shed light on novel approaches to repolarize GAMs to a more anti-tumorigenic state, as such presenting an alternative approach to ultimately improve treatment of GBM patients (Yang *et al*, 2023).

In recent years, the knowledge of GBM pathophysiology has advanced significantly and many promising research strategies have been pursued. The zebrafish (*Danio rerio*) xenograft model has proven to be a versatile animal model ideally suited for cancer research (Chen *et al*, 2021b). Furthermore, proof-of-concept studies have suggested that zebrafish xenograft models can serve as a preclinical drug screening platform and open the possibility to guide personalized treatments (Zon & Peterson, 2005; di Franco *et al*, 2022; Veinotte *et al*, 2014). Moreover, most relevant brain regions and the blood-brain barrier (BBB) are highly conserved and thus have a similar structure compared to humans. The high physiological conservation and the simplicity of xenotransplantation of human cancer cells into zebrafish embryos, have recently enabled researchers to recapitulate the characteristics of GBM, including the TME (Reimunde *et al*, 2021). However, studies that investigate the dynamic interactions between patient-derived GBM tumor cells and GAMs through long-term real-time recordings are currently lacking, though these insights are crucial to understand how GAMs influence disease progression.

In this work, we leverage a combination of single-cell RNA sequencing (scRNA-seq) of *in vitro* GAM-GBM co-cultures, real-time *in vivo* monitoring of GAM-GBM interactions in orthotopic zebrafish xenograft models to map cellular, molecular and spatial heterogeneity of GBM and associated macrophages. We extensively characterize a set of eight patient-derived GBM stem cell cultures (PD-GSCCs, hereafter referred to as GSCC in short) with different genomic and transcriptomic profiles, in an *in vitro* co-culture model with human macrophages. Using scRNA-seq, we report the heterogeneity in molecular changes in the macrophages influenced by the various GSCCs and identify patient-specific interaction patterns. We also show the use of high-resolution live-imaging in an orthotopic zebrafish xenograft model to visualize the dynamic interactions between transplanted tumor cells and GAMs in real-time. We developed an image analysis pipeline to process *in vivo* recordings, which was used to identify distinct behavioral patterns of GSCCs and associated macrophages. The time-lapse movies reveal tumor cell invasion and infiltration of reactive macrophages and how this varies across patients. Ultimately, using differential gene expression (DGE) analysis, immunohistochemistry (IHC) profiling of original tumor samples, and knock-out (KO) experiments, our work identifies Galectin-1 (GAL1, *LGALS1)* as an important immunomodulating target that affects tumor growth.

## Results

### Experimental set-up: studying the interaction between macrophages/GAMs and GSCCs in model systems

To analyze the cellular and molecular interaction of macrophages/GAMs and GSCCs, we made use of both an *in vitro* and *in vivo* model (Fig 1A). First, cellular interactions were studied in a co-culture model using scRNA-seq. Human monocytes were isolated from the blood of healthy volunteers, differentiated into macrophages, and co-cultured with eight different GSCCs, originating from seven different GBM patients. For one patient, we included two paired GSCCs (CME037/CME038), generated from a sample at initial diagnosis and recurrence. Importantly, the GSCCs used for these experiments cover a broad spectrum of genetic aberrations that are frequently present in GBM (Fig 1B): GSCCs were IDH-1-wildtype with mutations and/or copy number variations in common tumor suppressors and oncogenes such as EGFR^amp/mut^/Chr7^amp^, CDKN2A^del^, TP53^mut^, PTEN^mut/del^/Chr10^del^, CDK4^amp^, BRAF^amp/mut^ and NF1^mut^. We performed OneSeq analysis (Agilent) combining focused exome sequencing of the 43 most mutated genes in GBM with genome-wide copy number variation analysis of the GSCCs, and identified both shared and GSCC-specific mutations, consistent with previously reported intertumoral genomic heterogeneity of GBM (Rosenberg *et al*, 2017).

**Figure 1:**
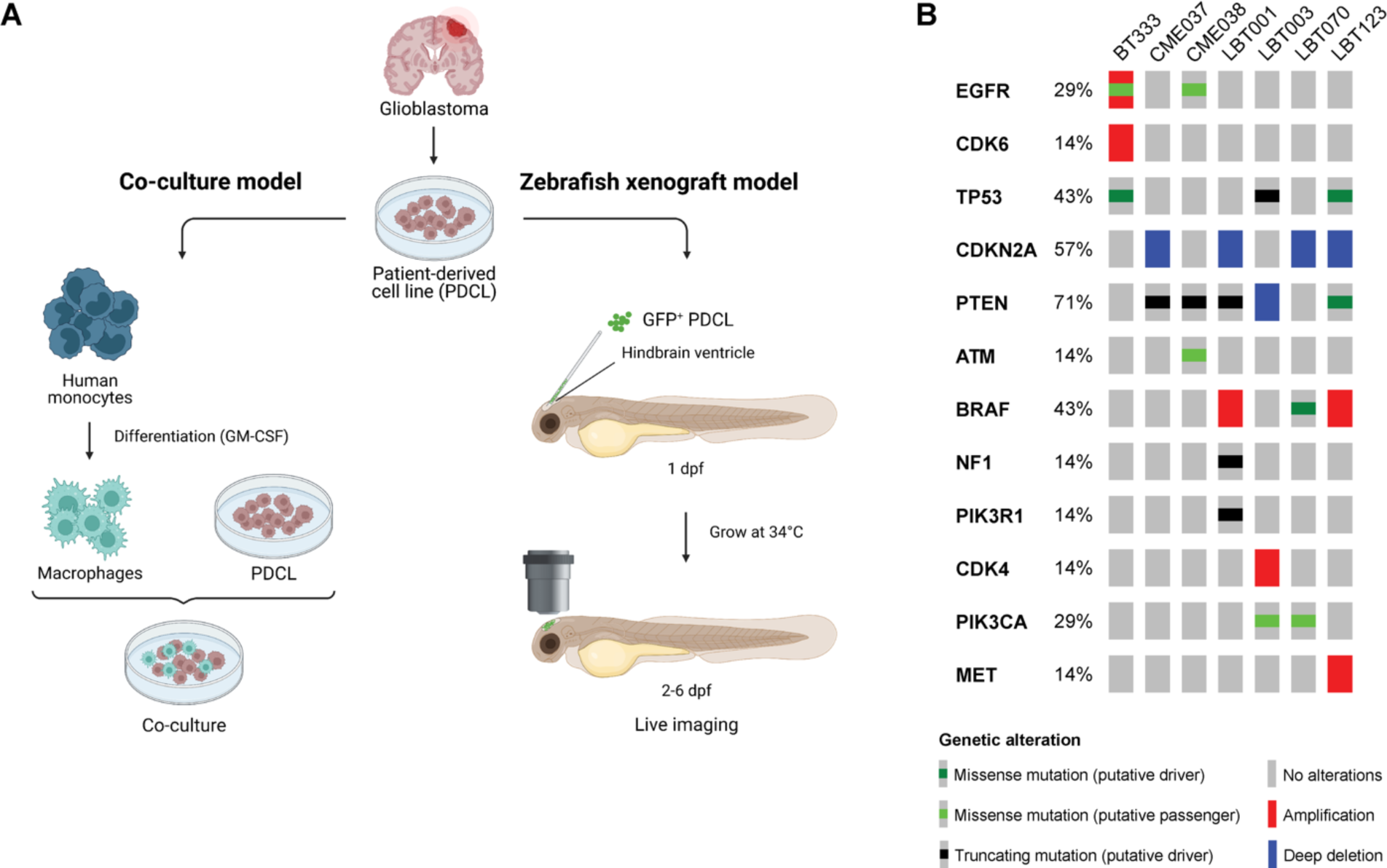
Overview experimental set-up and included GSCCs. A Schematic overview of the study design. Eight different GSCCs were used in a co-culture model with human monocyte-derived macrophages (left) and in an orthotopic zebrafish xenograft model (right). dpf = days post fertilization. B Oncoprint of included GSCCs: broad spectrum of common genetic aberrations in GBM were covered. Included GSCCs are mentioned on top of the figure.

Next, we optimized an orthotopic zebrafish xenograft model (hereafter referred to as zebrafish avatar model) in which GFP^+^ GSCCs were engrafted into a macrophage reporter line (see Materials and Methods, Fig 1A). This was achieved by injecting GFP^+^ GSCCs in the hindbrain ventricle of zebrafish embryos at 30 hours post fertilization (hpf). Zebrafish avatars were subsequently grown at an elevated temperature of 34°C (as compared to 28°C at which zebrafish are normally grown) to allow normal development of the embryos, while maintaining an environment that also supports tumor cell proliferation. Dynamic interactions between GAMs and GSCCs were eventually captured in time-lapse movies by performing live-imaging of the zebrafish embryos.

### Single-cell profiling of co-cultures of GSCCs and macrophages

To investigate how GSCCs influence the phenotypic features of macrophages in a patient-specific manner, we established an *in vitro* co-culture system between GSCCs and human monocyte-derived macrophages (Fig 2A). Macrophages and GSCCs were co-cultured in a 1:5 ratio using the hanging-drop method (Keller, 1995; Foty, 2011), while hanging drops of only macrophages were used as a control. After four days, cells of the various conditions were collected, dissociated, labeled using the MULTI-seq methodology (McGinnis *et al*, 2019) and pooled for single-cell RNA-seq. Subsequent sequencing and demultiplexing yielded a total of 5320 cells from 9 samples (8 co-cultures + 1 monoculture of macrophages, 218-1182 cells per sample) that passed quality control thresholds (see Materials and Methods), with a median of 3334 genes detected per cell (Fig S2A). Dimensionality reduction and unsupervised clustering (see Materials and Methods) revealed a clear separation between GSCCs and macrophages, with sample-specific clustering of the GSCCs and co-clustering of all macrophages (Fig 2C-D). As anticipated, the paired GSCCs (CME037/CME038) clustered together. Each cluster was identified as GSCCs or macrophages based on the expression of *SOX2* and *CD68* respectively (Fig S2B-C). Proliferating GBM tumor cells were identified based on their cell cycle score (Fig S2D).

**Figure 2:**
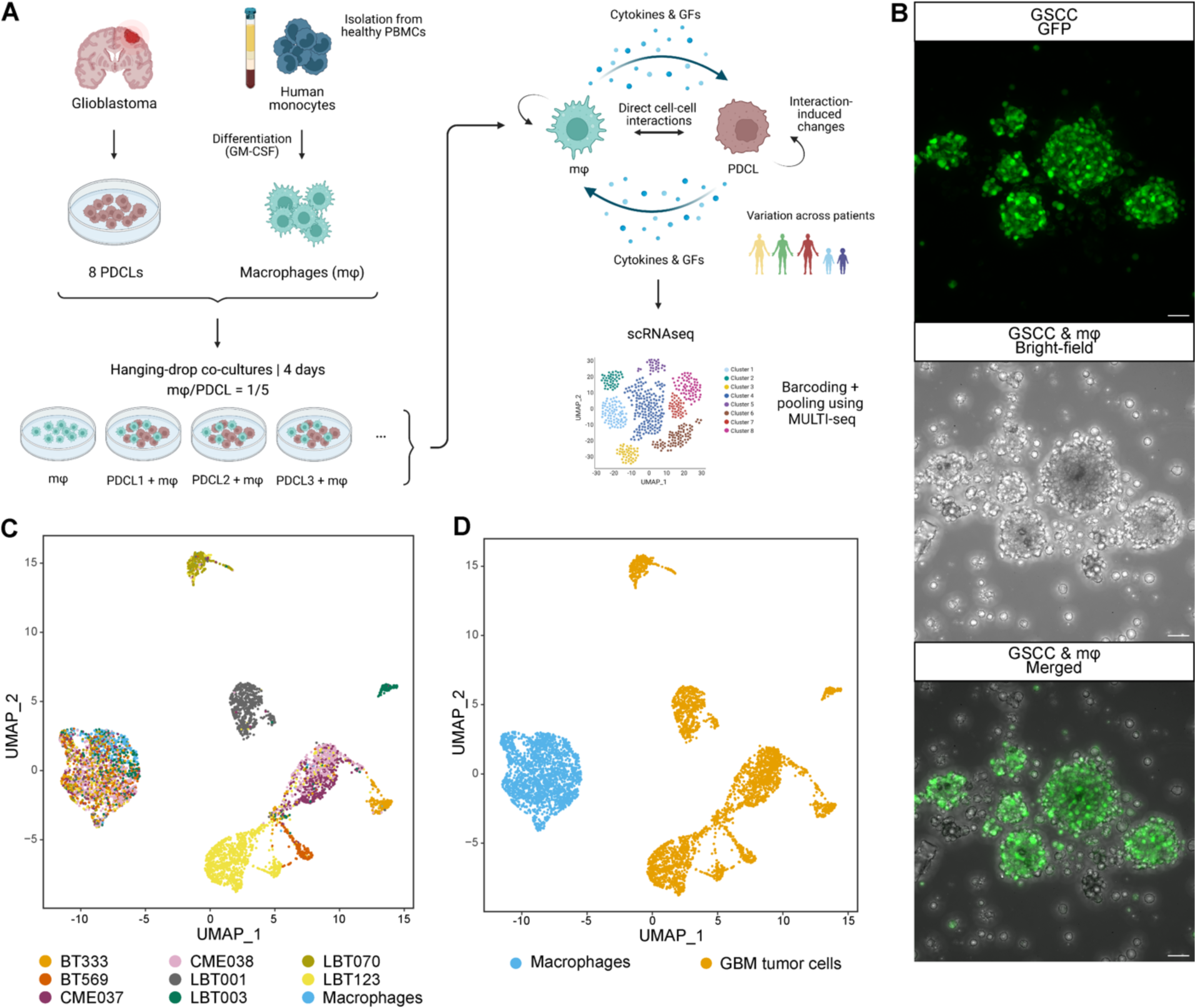
Single-cell profiling of GSCC-macrophage co-cultures. A Schematic overview of the experimental set-up of the scRNA-seq profiling assay. PBMCs, peripheral blood mononuclear cells; GFs, growth factors. B High-resolution images of a GSCC-macrophage co-culture. GFP^+^ GSCC was used for visualization of the co-culture with non-labeled macrophages. Scale bars: 50 μm. C-D Uniform Manifold Approximation and Projection (UMAP) plots of 5320 cells from 9 samples, annotated by sample name (C) and by cell type (D).

### Molecular heterogeneity of GBM-associated macrophages

GAMs can represent up to 50% of GBM tumors. To elucidate the molecular heterogeneity of the monocyte-derived macrophages in the co-culture model, we performed unsupervised subclustering of the macrophages, yielding three clusters (MC1-3) (Fig 3A + S3A). All GSCCs contributed to each macrophage cluster (Fig S3B). As reported by others, the observed macrophage subtypes did not fit the classical, yet outdated, M1/M2 macrophage paradigm, introduced by Mills and colleagues (Mills *et al*, 2000; Müller *et al*, 2017). There was limited analogy between well-known M1 and M2 markers and macrophage cluster signature genes. However, the expression levels of well-known markers for immune-stimulation and -suppression were investigated, where we found that MC1 macrophages had a more immunosuppressive nature, while MC2 macrophages exhibited more immune-stimulatory features (Fig 3B).

**Figure 3:**
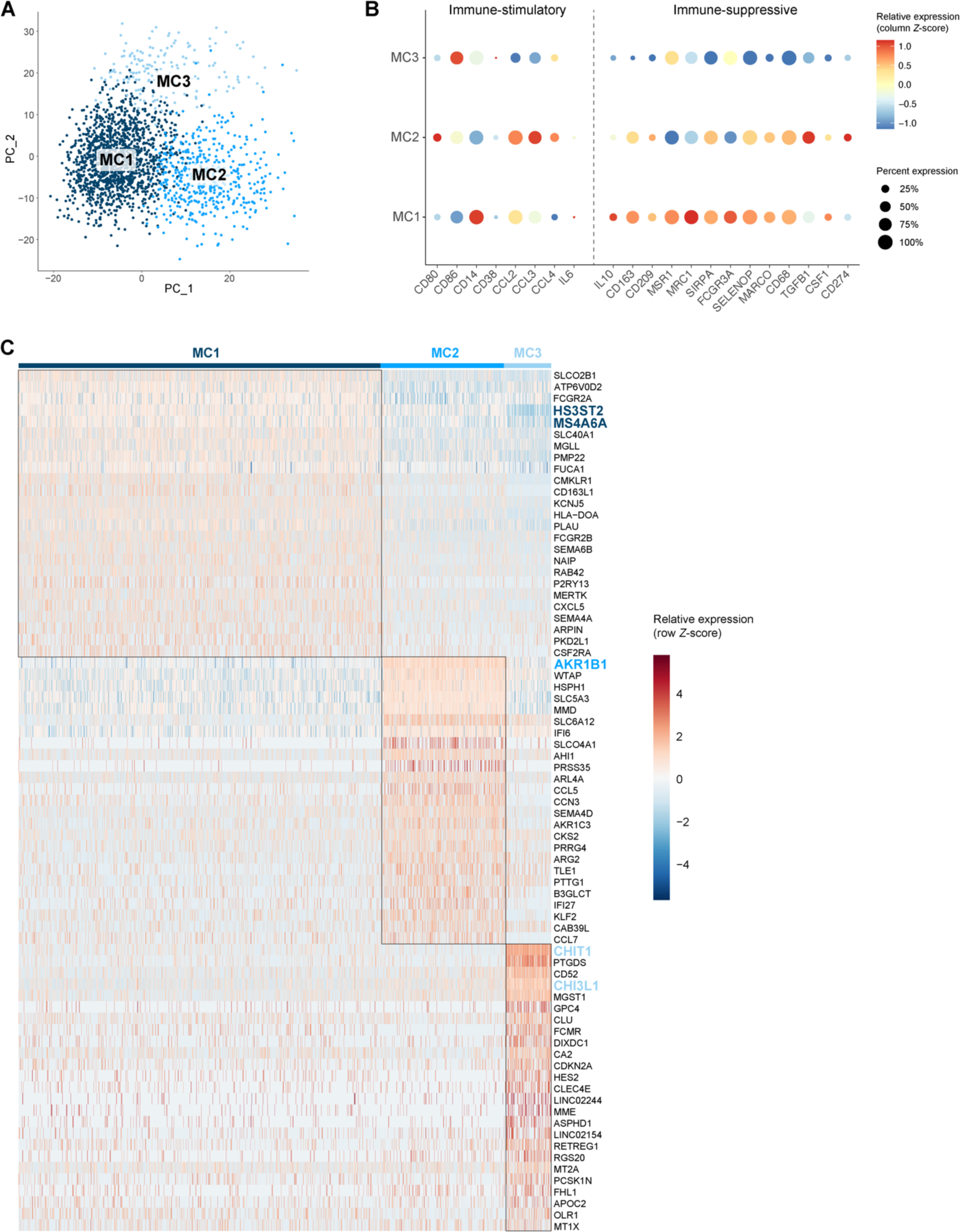
Molecular heterogeneity of GBM-associated macrophages. A Principal component analysis (PCA) plot of macrophage population identified three distinct macrophage subclusters (MC1-3). PCs were calculated using the 2000 most variable genes. Plot shows PC1 and PC2. B Dot plot showing marker gene expression for immune-stimulation and -suppression. Dot size indicates the percentage of cells in each macrophage subcluster expressing the gene, and dot color indicates the relative expression level. C Heatmap of top 25 differentially expressed genes in the macrophage subclusters, ranked by log2(FC). Genes discussed in the text are highlighted in the subcluster colors.

Next, we manually annotated the distinct macrophage subtypes using marker gene analysis (Fig 3C). MC1 consisted of macrophages that expressed high levels of immunosuppressive genes *HS3ST2* and *MS4A6A*. *HS3ST2* has previously been shown to be highly expressed upon alternative stimulation (M2 polarization), while the enzyme was not detected in pro-inflammatory macrophages and primary monocytes (Martinez *et al*, 2015). Interestingly, *MS4A6A* is expressed in M2 macrophages and correlates with macrophage infiltration, unfavorable clinical outcome, and poor responses to adjuvant chemotherapy in glioma patients (Zhang *et al*, 2022). MC2 contained macrophages that express *AKR1B1* (Fig 3C), which is commonly associated with an inflammatory profile (Erbel *et al*, 2016; Cheng *et al*, 2021). Of note, cells in MC2 mainly originated from the macrophage monoculture and represent cells that were polarized towards an inflammatory subtype induced by treatment with granulocyte-macrophage colony-stimulating factor (GM-CSF) (Lotfi *et al*, 2020). MC3 macrophages expressed high levels of *CHIT1* (and *CHI3L1*) and lower levels of the mature macrophage markers *CD68*, *CD163*, *CD204/MSR1*, and *CD206/MRC1* (Fig 3B-C). As such, we identified MC3 as transitioning monocytes (TransMos) as it has been demonstrated that *CHIT1* expression increases exponentially over time in monocytes during the transition to macrophages (di Rosa *et al*, 2013).

### Macrophages shift towards an immunosuppressive phenotype upon co-culture with patient-derived GSCCs

We then sought to characterize the molecular switch in macrophages induced by interactions with GSCCs by performing pseudotime analysis. MC3 was not included in the analysis because we aimed to capture the interaction between mature macrophages and GSCCs, rather than the interaction between TransMos and GSCCs (Fig S4A). The bulk of the variance between MC1 and MC2 is explained by principal component (PC) 1 (Fig 4A). The macrophage subcluster distribution across the GSCCs indicated that macrophages switch to an immunosuppressive state upon co-culture with GSCCs (MC2 to MC1, Fig 4B). The macrophage monoculture mainly consisted of inflammatory MC2 macrophages, induced by treatment with GM-CSF. Upon co-culturing, the proportion of MC2 macrophages significantly decreased for all GSCCs (Chi-square test, *p* < 0.0001). Surprisingly, co-cultures of LBT003 contained significantly more pro-inflammatory MC2 macrophages and less immunosuppressive MC1 macrophages than co-cultures of other GSCCs (Chi-square test, *p* < 0.0001) (Fig 4B).

**Figure 4:**
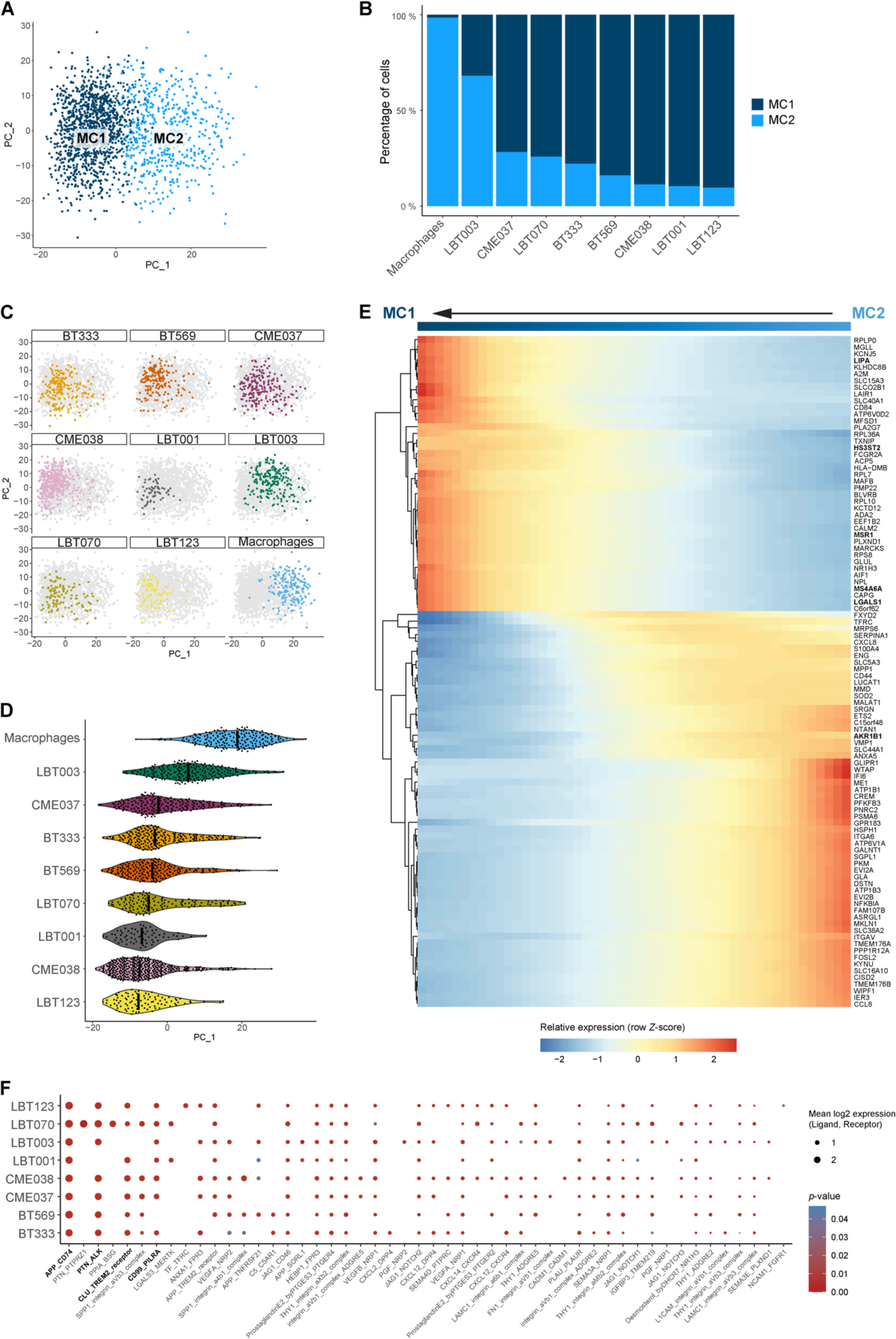
Macrophages shift towards an immunosuppressive phenotype upon co-culture with patient-derived GSCCs. A PCA plot of macrophage population without TransMos shows two distinct macrophage subclusters (MC1-2). PCs were calculated using the 2000 most variable genes. PC1 and PC2 are shown on the PCA plot. B Macrophage subcluster distribution for the different samples. C Representation of original samples on PCA plot. D Pseudotime analysis along PC1 axis. Violin plots depicting the PC1 values of each single cell, split up by sample. E Heatmap of the 100 most significant temporally expressed genes along the PC1 axis, constructed by fitting a generalized additive model with the PC1 value as a LOESS term (see Materials and Methods). F Dot plot of cell–cell communication analysis using CellPhoneDB. Depicted are L:R pairs for GSCC – macrophage signaling across all GSCCs, ranked by mean log2 expression. Each dot size shows the log2 mean of expression values and dot color indicates the *p*-value for the listed L:R pairs (x-axis) in the respective GSCCs (y-axis). Only top 50 significant L:R pairs, with cut-offs of *p*-value ≦ 0.05 are shown. The *p*-values were generated by CellPhoneDB, which uses a one-sided permutation test to compute significant interactions.

Next, we used each cell’s PC1 value as a measure for pseudotime along the MC2-MC1 trajectory and considered MC2 the root of the trajectory (Fig 4C-D). Principal component analysis (PCA) and Uniform Manifold Approximation and Projection (UMAP) plots per sample showed a gradient in the macrophage population (Fig 4C + S4B), indicating that macrophages were in the process of a phenotypic switch. Interestingly, of all samples, macrophages co-cultured with LBT003 most closely resembled the control macrophages (Fig 4C-D). To study which genes were involved in the phenotypic shift, we identified temporally expressed genes along the PC1 axis (from MC2 to MC1) by fitting a generalized additive model with the PC1 value as a LOESS term (see Materials & Methods, Fig 4E). We found that the expression of immunosuppressive genes increased along the MC2-MC1 trajectory, while the expression of immunostimulatory genes gradually decreased, thereby triggering a phenotypic change. For instance, immunostimulatory genes like *AKR1B1* and *CCL4* gradually decreased in expression along the trajectory (Fig S4C-D). *MSR1, LIPA* and *LGALS1,* genes that have previously been shown to be upregulated in immunosuppressive macrophages (Huang *et al*, 2014), were initially downregulated and increased in expression along the trajectory (Fig S4E-G).

To investigate whether potential ligand:receptor (L:R) interactions between GSCCs and macrophages could explain the observed macrophage polarization, we performed CellPhoneDB analysis (Fig 4F) (Garcia-Alonso *et al*, 2022). This revealed several strong putative interactions between GBM tumor cell ligands and macrophage receptors that were conserved over the different co-cultures, including APP:CD74, CD99:PILRA, PTN:ALK and CLU:TREM2. CD74 signaling, which was found to be significant in all co-cultures, has previously been shown to induce an inflammatory macrophage phenotype (Zeiner *et al*, 2015). In contrast, TREM2 signaling induces a strong immunosuppressive phenotype and elevated TREM2 expression in macrophages is considered a bad prognostic marker in several pathologies (Katzenelenbogen *et al*, 2020; Yu *et al*, 2023). Interestingly, putative CLU:TREM2 signaling was not significant for LBT003, which induced the weakest macrophage polarization in the co-culture model, suggesting that the lack of this interaction might in part explain the limited polarization. When investigating predicted macrophage to GSCC signaling, several conserved putative interactions were found, including APLP2:PLXNA4, LRPAP1:SORT1, MDK:PTPRZ1, SIRPA:CD47 and TYROBP:CD44 (Fig S4H). Of note, CD47, which is known to be upregulated in glioma stem cells, binds to SIRPα on the surface of macrophages to exert a “don’t eat me” signal, thus contributing to immune evasion of the cancer cells by preventing phagocytosis by macrophages (Hu *et al*, 2020; Gholamin *et al*, 2017).

In general, we found that the interaction between GSCCs and GAMs induced a shift to a more immunosuppressive phenotype, but the magnitude of this shift was variable across GSCCs. Strikingly, CME037 and CME038, which are derived from the same patient at initial diagnosis and recurrence respectively, were positioned at opposite ends of the spectrum, with the recurrent GSCC exhibiting a more immunosuppressive polarization. Furthermore, we showed multiple predicted interactions between GSCCs and macrophages that could govern macrophage polarization or immune evasion, some of which were conserved over all co-cultures, while others were specific to certain cultures.

### Mapping the real-time evolution of GSCCs and associated macrophages in zebrafish avatars

To investigate the behavior of different GSCCs *in vivo*, we generated an orthotopic xenograft model in zebrafish embryos. The eight GSCCs that were used in the co-culture assay were stably labeled with GFP using viral transduction and injected into *Tg(mpeg1:mCherryF)^ump2^; Tg(kdrl:lynEYFP)* zebrafish embryos, characterized by mCherry-expressing macrophages (and microglia) and YFP-labeled vasculature, at 30 hpf (Ellett *et al*, 2011; Silva *et al*, 2021). Next, live-imaging of these zebrafish avatars was performed to directly capture dynamic interactions between xenografted GSCCs and macrophages. Zebrafish avatars were typically imaged for 8-16 hours at two different timepoints to follow tumor progression: at 1 day post injection (dpi) and again at 5 dpi (Fig 5A + S5A). Based on the morphology of the macrophages, a distinction was made between round and ramified macrophages (Fig 5B). We developed an image analysis pipeline for image processing in 3D and over time to compute GSCC-specific morphometrics and dynamics of the tumor and its microenvironment (Fig 6A).

**Figure 5:**
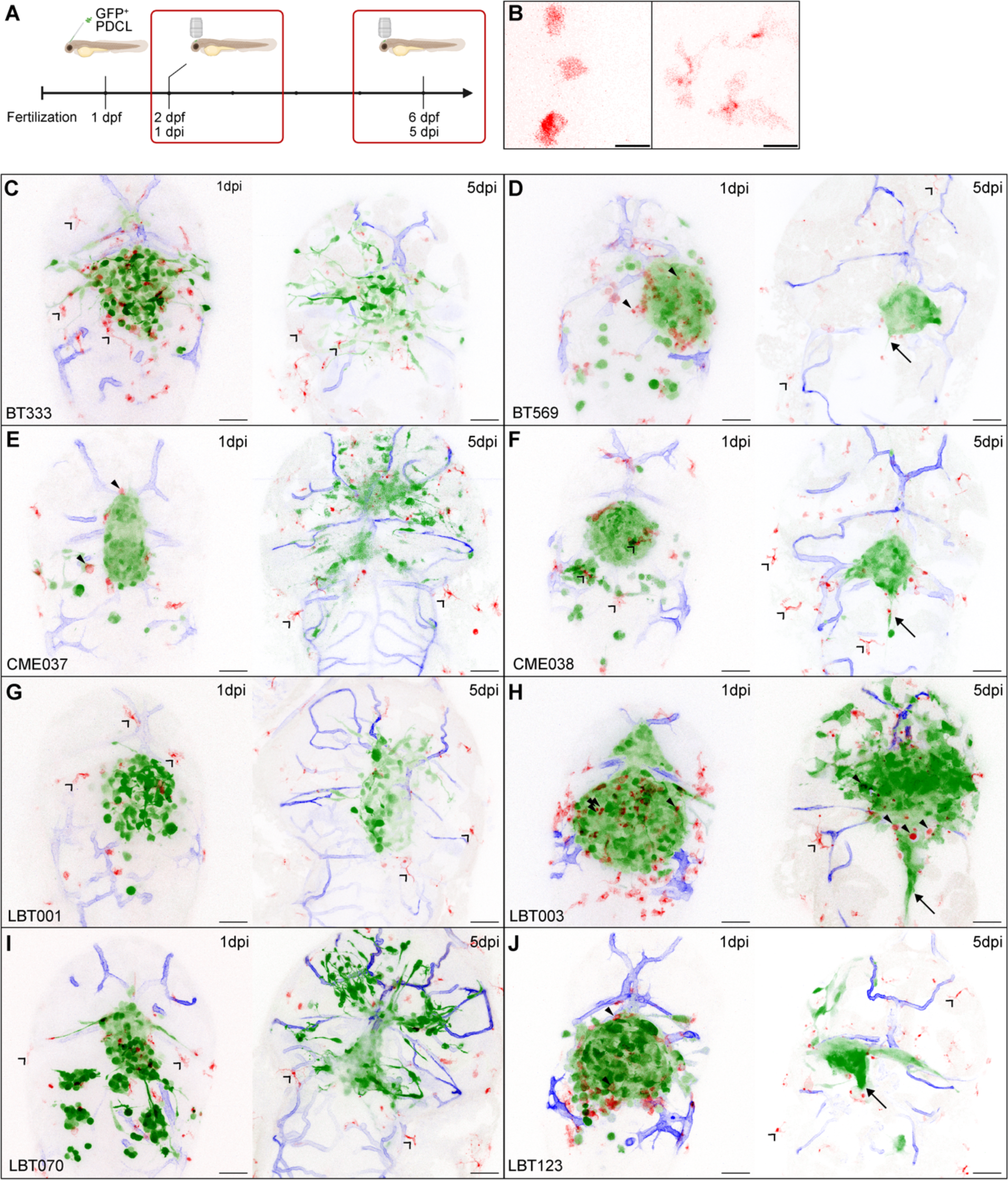
Overview of different zebrafish avatars. A Schematic overview showing the timeline of the orthotopic zebrafish xenograft model. B Zoomed-in images of round (left) and ramified (right) macrophages in the recorded time-lapse movies. Scale bars: 15 μm. C-J Representative maximum-projections of a z stack of the head region of *Tg(mpeg1:mCherryF)^ump2^; Tg(kdrl:lynEYFP)* zebrafish embryos with different GFP-labeled patient-derived GSCC tumors, at 1 dpi (left panel) and 5 dpi (right panel). (C) BT333; (D) BT569; (E) CME037; (F) CME038; (G) LBT001; (H) LBT003; (I) LBT070; (J) LBT123. GBM tumor cells are shown in green, macrophages in red, and blood vessels in blue. Arrows indicate midline protrusion of the tumor. Filled arrowheads indicate round GAMs. Open arrowheads indicate ramified GAMs. Scale bars: 50 μm.

**Figure 6:**
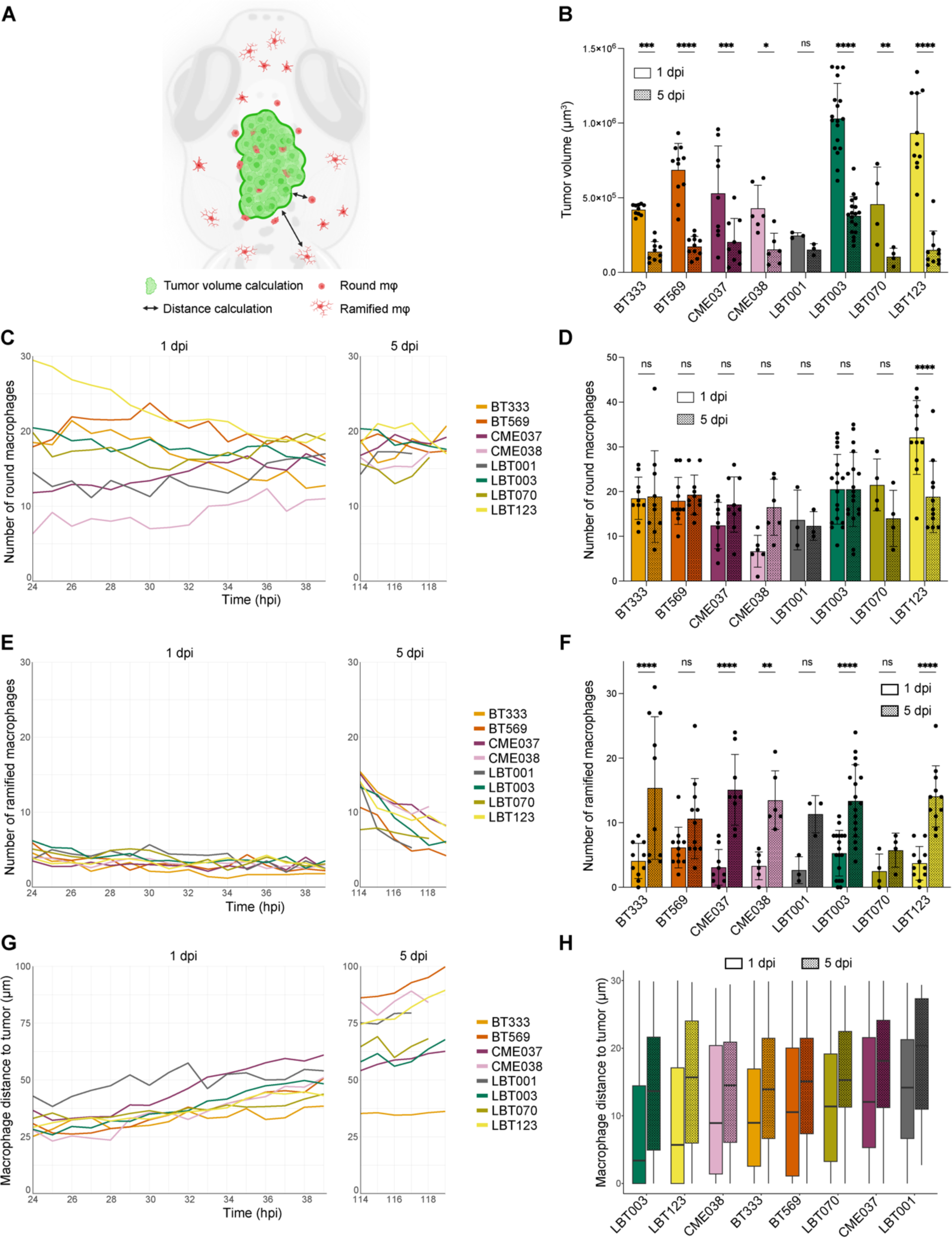
GSCC-specific morphometrics and dynamics of the tumor and its microenvironment in 3D over time. A Schematic overview of features extracted and computed from the time-lapse movies of the orthotopic zebrafish xenograft model. B Tumor volume at the start of 1 dpi and 5 dpi time-lapse movies. (*n* = 3-18 zebrafish embryos per group, with an average of 9 embryos per group). C Mean number of round macrophages over time, during 1 dpi movies (left) and 5 dpi movies (right). (*n* = 4-22 zebrafish embryos per group, with an average of 10.81 embryos per group). D Number of round macrophages at the start of 1 dpi and 5 dpi time-lapse movies. (*n* = 3-18 zebrafish embryos per group, with an average of 9 embryos per group). E Mean number of ramified macrophages over time, during 1 dpi movies (left) and 5 dpi movies (right). (*n* = 4-22 zebrafish embryos per group, with an average of 10.81 embryos per group). F Number of ramified macrophages at the start of 1 dpi and 5 dpi time-lapse movies. (*n* = 3-18 zebrafish embryos per group, with an average of 9 embryos per group). G Mean macrophage distance to the tumor over time, during 1 dpi movies (left) and 5 dpi movies (right). (*n* = 107-945 macrophages per group, with an average of 455.31 macrophages per group). H Boxplot of macrophage distance to the tumor of all macrophages within 30 μm of the tumor, at the start of 1 dpi and 5 dpi time-lapse movies, ranked by increasing median distance at 1 dpi. (*n* = 26-634 macrophages per group, with an average of 208 macrophages per group; boxes stand for 50% of the data and minima/maxima are indicated by the line ends). Data information: In B+D+F, data are presented as mean ± SD. The *p*-values were calculated by two-way repeated measures ANOVA, followed by Šidák’s multiple comparisons correction. ns ≥ 0.05, \**p* < 0.05; \*\**p* < 0.01; \*\*\**p* < 0.001, \*\*\*\**p* < 0.0001.

Although for most GSCCs the tumor tended to decrease in size, we observed a clear difference in the way tumors evolved over time (Fig 5C-J + Movie 1-16). Some GSCCs (BT333, CME037, LBT001 and LBT070) were more invasive, while other GSCCs (BT569, CME038, LBT003 and LBT123) were more confined. Cell numbers of injected GSCCs typically declined from 1 dpi onwards, with significant reduction to about 30% of the original tumor volume at 5 dpi for all GSCCs, except for LBT001 (Fig 6B + S6A), suggesting that cancer cell proliferation was slower than clearance by the embryos’ immune system. Tumors of BT333, CME037, LBT001, and LBT070 showed a more invasive phenotype with infiltrating cells and shape changes over time (Fig 5C+E+G+I). In general, GSCCs displayed invasive protrusions at 1 dpi, and their number even increased towards 5 dpi. Other tumors like BT569, CME038 and LBT123 preferred to establish compact and round tumor masses, while forming a protrusion along the midline (Fig 5D+F+J, indicated with arrows). Interestingly, LBT003 tumors were significantly larger than most other tumors at 1 dpi, and this was even more pronounced at 5 dpi (Fig 5H + S6B-C). LBT003 tumors also formed a protrusion along the midline (Fig 5H, arrow). Furthermore, we noticed two different trends in the 1 dpi movies: tumors that decreased in size (BT333, CME037, CME038, LBT003, LBT070 and LBT123), and tumors that slightly grew over time (BT569 and LBT001), while from 5 dpi onwards, tumor size remained roughly unchanged (Fig S6A).

### Live recordings of dynamic macrophage interactions reveal higher infiltration of reactive macrophages in LBT003 avatars

Next, we assessed the interaction between GAMs and engrafted GSCCs over time. We observed two types of macrophages: round macrophages that were compact in shape and located in proximity of the tumor, and ramified macrophages characterized by an elongated cell body and formation of filopodia-like structures that were located further away from the tumor (Fig 5B+C-J). These morphological phenotypes were described previously and correlate to different functions: round macrophages exert antitumoral properties, while ramified macrophages are more homeostatic and immunosuppressive (McWhorter *et al*, 2013; Heindl *et al*, 2018). Interestingly, our high-resolution time-lapse recordings revealed close interaction between round macrophages and GBM tumor cells. Round macrophages appeared to attack and even phagocytize GBM tumor cells, shown by co-localization of GBM tumor cells and macrophages (Fig S5B + Movie 17).

Generally, BT569, CME037, LBT123 and especially LBT003 tumors attracted a lot of round macrophages at 1 dpi (Fig 5D+E+H+J, filled arrowheads), while less macrophages were present around BT333, CME038, LBT001, and LBT070 tumors (Fig 5C+F+G+I + Movie 1-16). Nevertheless, some of the latter GSCCs attracted some ramified macrophages (Fig 5C+F+G+I, open arrowheads). At 5 dpi, we observed few macrophages overall, which were mostly ramified and did not interact with the GSCCs (Fig 5C-J, open arrowheads). LBT003 avatars were an exception, as round macrophages were present close to the tumor even at 5 dpi (Fig 5H, filled arrowheads).

Quantitative analysis demonstrated that the number of round macrophages decreased for most GSCCs during the time-lapses starting at 1 dpi, while the number of ramified macrophages remained the same (Fig 6C-F). However, for the paired GSCCs (CME037/CME038) and LBT001, the number of round macrophages slightly increased over time in 1 dpi movies (Fig 6C). Remarkably, at the start of 5 dpi time-lapses the number of ramified macrophages was significantly higher than at 1 dpi for most GSCCs (Fig 6E-F), while the number of round macrophages was roughly the same at 1 and 5 dpi (Fig 6C-D). This suggests that macrophages switch towards an immunosuppressive phenotype over time.

Next, we calculated the three-dimensional distance of the macrophages to the tumor, and found that in general the distance of the macrophages significantly increased during the 1 dpi movies (*p* = 5.96e-7), while this trend was not significant in the 5 dpi movies (*p* = 0.71) (Fig 6FG+ S6D-E). Furthermore, for all GSCCs, the round macrophages were located significantly closer to the tumor than the ramified macrophages (*p* < 2.2e-16) (Fig S6F), indicating that the round phenotype correlates with reactive and antitumoral properties. Finally, we considered only the macrophages within 30 µm of the tumor, as those are the ones interacting most closely with the tumor. We found that within this distance, macrophages were located closer to LBT003 tumors at 1 dpi and 5 dpi, compared to other tumors, although less pronounced at 5 dpi (Fig 6H + S6G-H).

Taken together, these observations illustrate that LBT003 avatars have higher infiltration of reactive macrophages compared to other GSCCs.

### Macrophage/GAM - GSCC interactions correlate with clinical outcome in GBM patients

Since both the co-culture model and the zebrafish avatar model identified LBT003 as distinct from the other GSCCs, we were interested in the clinical background of this patient. Interestingly, this patient was diagnosed in the beginning of 2017 with GBM, received SoC treatment, and is still alive today (Fig 7A). With a survival of 71 months (and still counting) this is 4.7 times longer than the median survival of a GBM patient receiving SoC treatment (*i.e.* 15 months) (Grochans *et al*, 2022), and this patient can be considered a “long-term survivor” (Decraene *et al*, 2023). This suggests that this patient is not only an outlier at the molecular level, as our models have demonstrated, but also in a clinical context. Further research is required to discern whether one is causative of the other.

**Figure 7:**
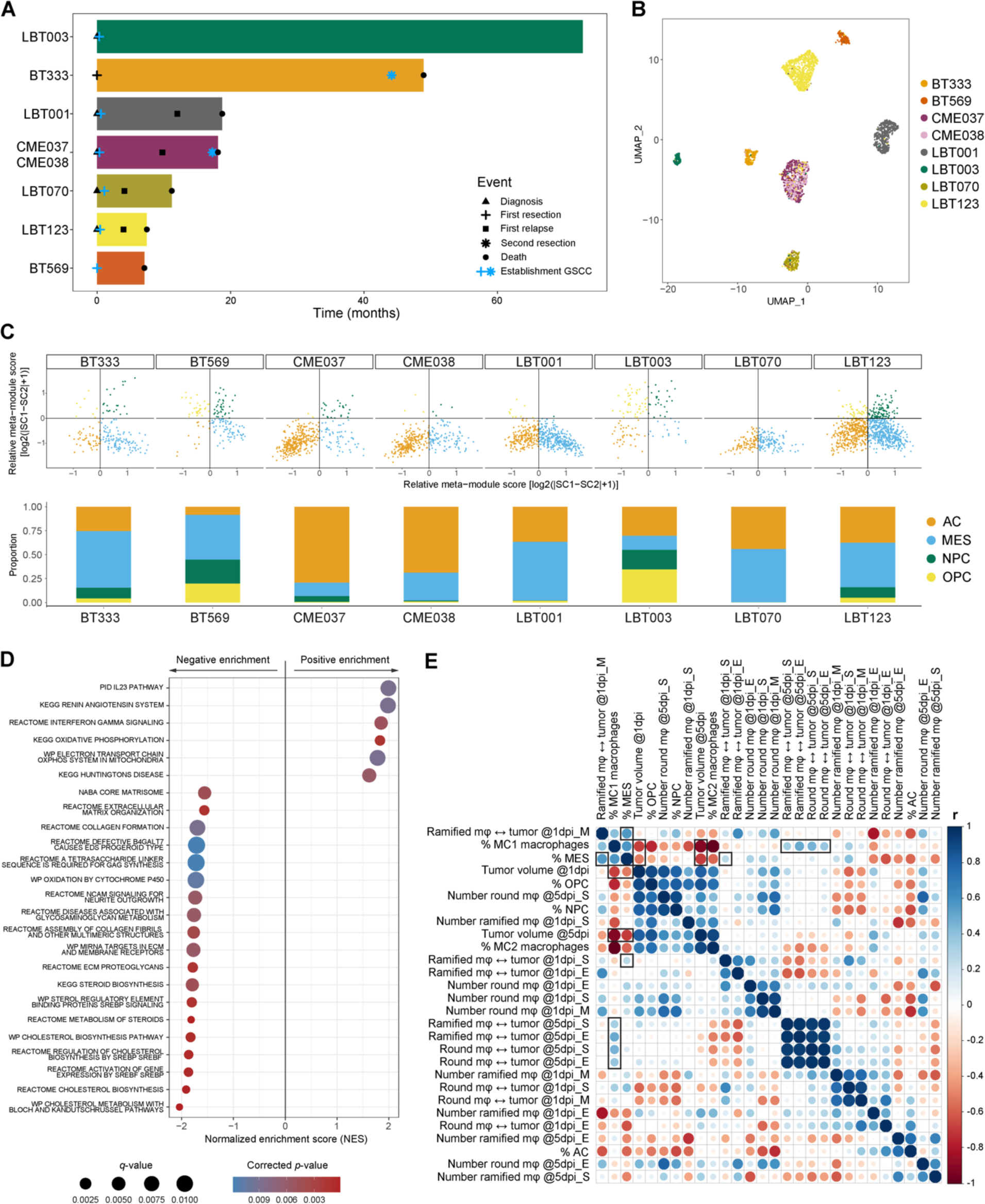
Macrophage/GAM-GSCC interactions correlate to clinical outcome in GBM patients. A Swimmer plot of included patients with indication of important events. B UMAP plot of GBM tumor cells annotated by sample name. C Two-dimensional butterfly plot visualization of molecular subtype signature scores according to Neftel *et al*. (top). Each quadrant corresponds to a subtype, and the position of each cell reflects its relative signature scores. Colors represent different clusters. Neftel cluster distribution for the different samples (bottom). D Dot plot showing the 25 most significant positive and negative enriched GSEA pathways using all curated gene sets of WikiPathways, Reactome, KEGG, PID and BioCarta databases for the group of invasive GSCCs (cut-off corrected *p*-value = 0.05). E Correlogram of *in vitro* and *in vivo* data of the GSCCs (*n* = 8). Dot size and color indicate the Pearson correlation coefficient (r). mφ, macrophages; ↔, distance to; S, start (24-26 or 114-116 hpi); M, mid (30-32 hpi); E, end (37-39 or 117-119 hpi) indicate timepoints of the time-lapse movies.

Next, we performed unsupervised subclustering of the GBM tumor cells, which again revealed patient-specific clustering (Fig 7B + S7A). Neftel *et al*. previously reported that GBM tumor cells can be classified into oligodendrocyte-progenitor (OPC), neural-progenitor (NPC), astrocyte (AC), or mesenchymal (MES)-like cell states (Neftel *et al*, 2019). Stratifying single GBM tumor cells according to the gene sets described by Neftel *et al*., we found that most GSCCs consisted of a mix of all four subtypes, though at different relative proportions, further illustrating the intra- and intertumoral heterogeneity captured within these cell models (Fig 7C + S7B). Notably, LBT003 contained the smallest proportion of MES-like cells, which have previously been shown to be more invasive and to be associated with a worse prognosis (Stead, 2022), and is in line with the clinical evolution of LBT003. Interestingly, there was an increase in MES-like cells in CME038, the recurrent GSCC of CME037, consistent with a recently published study by Wang *et al*. (Wang et al, 2022b).

As we were interested in the overlap of our findings from the scRNA-seq data and the features extracted from the zebrafish avatar model, we first examined whether the transcriptomic profiles of the GSCCs could explain the differences in invasiveness observed in the avatar model. We previously observed two distinct patterns of tumor evolution in the zebrafish avatar model: invasive (BT333, CME037, LBT001 and LBT070) and non-invasive (BT569, CME038, LBT003 and LBT123). Therefore, we performed DGE analysis between the invasive and the non-invasive GSCCs on the scRNA-seq data from the co-culture experiments. Gene set enrichment analysis (GSEA) revealed an enrichment for oxidative phosphorylation (OXPHOS) in the invasive GSCCs and an enrichment for extracellular matrix (ECM) production and deposition in the non-invasive GSCCs (Fig 7D).

Next, we performed a more general correlation analysis to correlate the features extracted from both model systems (Fig 7E). The percentage of immunosuppressive MC1 macrophages was correlated with the distance of macrophages to the tumor at 5 dpi, indicating that immunosuppressive GSCCs were also associated with surveilling macrophages *in vivo*. In addition, the percentage of MC1 macrophages was negatively correlated with the tumor volume at 1 dpi and 5 dpi. Although this may seem contradictory, tumor size is not always related to invasiveness, aggressiveness and thus survival (Wang *et al*, 2019). A large, but well-aligned tumor, which is the case for LBT003 tumors, improves surgical resection and thus patient’s survival, compared to smaller invasive tumors (Mair *et al*, 2018). Finally, the percentage of MES-like cells was correlated with the distance of ramified macrophages at 1 dpi, and negatively correlated with the tumor volume at 1 and 5 dpi, which is in line with our finding that in LBT003 tumors, which contained the smallest proportion of MES-like cells, macrophages were located close to the tumor, and LBT003 tumors were bigger than other tumors.

#### *LGALS1* IS INVOLVED IN SUPPRESSION OF THE IMMUNE SYSTEM

To investigate whether the phenotypic differences between LBT003 and the other GSCCs can be explained by transcriptomic differences, we performed DGE analysis between GBM tumor cells from LBT003 versus the other GSCCs. We found *LGALS1* to be significantly downregulated in LBT003 (Fig 8A-B). Interestingly, *LGALS1* has been shown to be highly expressed in GBM tumor cells and drives therapy resistance (Rorive *et al*, 2001; Chou *et al*, 2018). While GAL1 is also present in the tumor stroma (Chou *et al*, 2018), research has indicated that tumor-derived, rather than TME-derived GAL1, is involved in the aggressiveness of glioma progression (Verschuere *et al*, 2014). Moreover, *LGALS1* silencing in GBM reduced macrophages’ polarization switch to an immunosuppressive state and was associated with increased survival (Van Woensel *et al*, 2017). The lower level of *LGALS1* expression in LBT003 could thus, at least in part, explain the phenotypic differences between LBT003 and the other GSCCs in the zebrafish avatar model (more GAMs situated closer to the tumor), as well as the general lack of macrophage polarization in the co-culture model.

**Figure 8:**
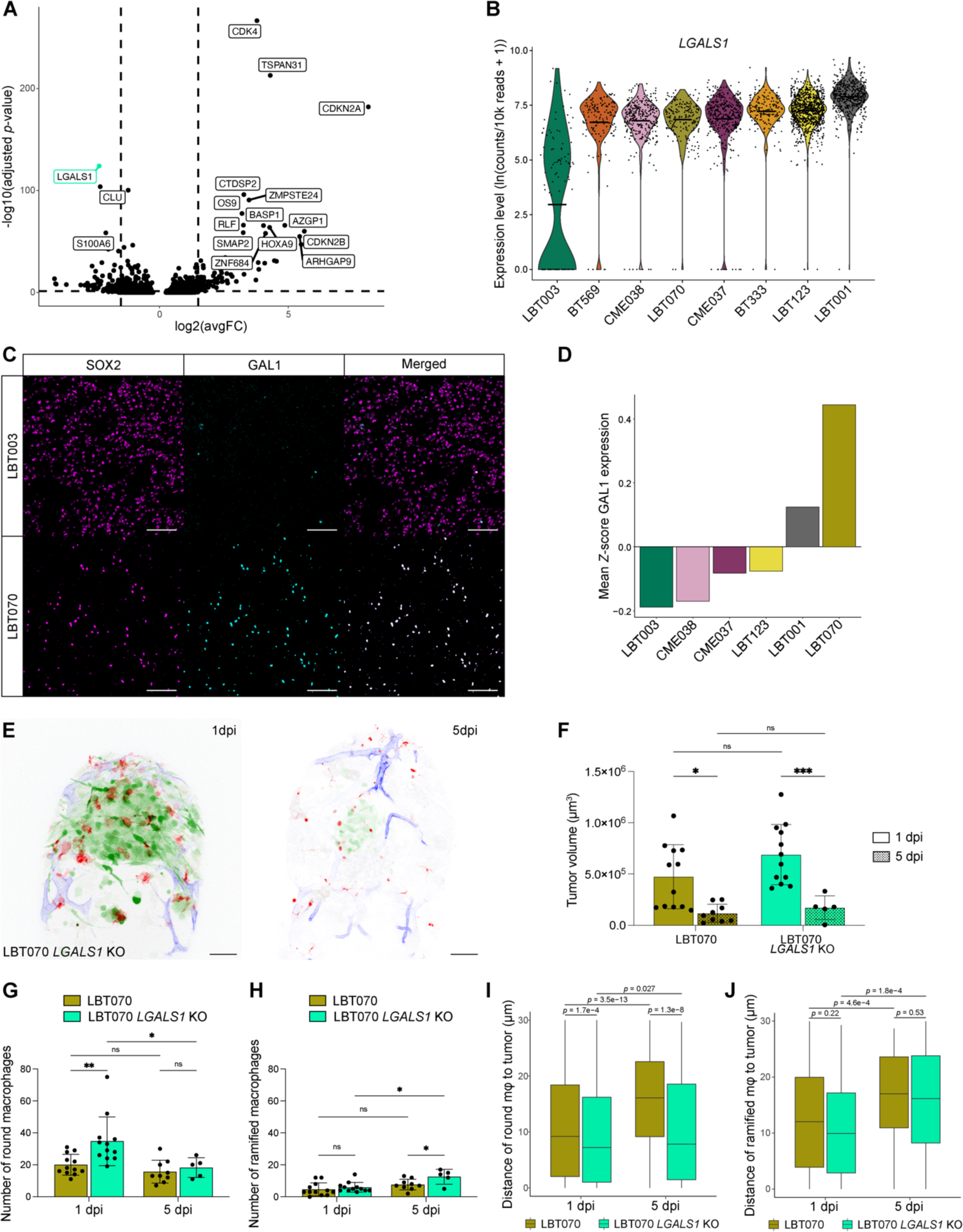
*LGALS1* is involved in suppression of the immune system. A Volcano plot depicting differentially expressed genes in LBT003 and all other GSCCs (Left: downregulated genes in LBT003; right: upregulated genes in LBT003.) Y-axis denotes -log10(adjusted *p*-value) while x-axis shows log2(avgFC) values. Cut-offs were set at -log10(adjusted *p*-value) > 1.3 and abs(log2(avgFC)) > 1.5. B Violin plot showing *LGALS1* expression levels in GSCCs. C Representative double immunofluorescence images showing co-expression of SOX2 (magenta) and GAL1 (cyan) in GBM tissue from LBT003 and LBT070. For enhanced visualization, a binary mask was generated from the SOX2^+^ cells and multiplied with the image of GAL1 staining in Fiji to exclude GAL1 staining in non-tumor cells. Scale bars: 100 μm. D Mean fluorescence intensity values for GAL1 staining in SOX2^+^ cells in GBM tissue samples, normalized using *Z*-scores within each sample. *n* = 6 tumor samples derived from 5 different patients. E Representative maximum-projections of a z stack of the head region of a *Tg(mpeg1:mCherryF)^ump2^; Tg(kdrl:lynEYFP)* zebrafish embryo with a GFP-labeled LBT070 *LGALS1* KO tumor, at 1 dpi (left panel) and 5 dpi (right panel). GBM tumor cells are shown in green, macrophages in red, and blood vessels in blue. Scale bars: 50 μm. F Tumor volume at the start of 1 dpi and 5 dpi time-lapse movies. (*n* = 5-12 zebrafish embryos per group, with an average of 9.5 embryos per group). G Number of round macrophages at the start of 1 dpi and 5 dpi time-lapse movies. (*n* = 5-12 zebrafish embryos per group, with an average of 9.5 embryos per group). H Number of ramified macrophages at the start of 1 dpi and 5 dpi time-lapse movies. (*n* = 5-12 zebrafish embryos per group, with an average of 9.5 embryos per group). I Boxplot of macrophage distance to the tumor of round macrophages within 30 μm of the tumor, at the start of 1 dpi and 5 dpi time-lapse movies. (*n* = 39-290 macrophages per group, with an average of 128.5 macrophages per group; boxes stand for 50% of the data and minima/maxima are indicated by the line ends). J Boxplot of macrophage distance to the tumor of ramified macrophages within 30 μm of the tumor, at the start of 1 dpi and 5 dpi time-lapse movies. (*n* = 13-40 macrophages per group, with an average of 25.5 macrophages per group; boxes stand for 50% of the data and minima/maxima are indicated by the line ends). Data information: In F-H, data are presented as mean ± SD, and the *p*-values were calculated by mixed-effects model, followed by Šidák’s multiple comparisons correction. ns ≥ 0.05, \**p* < 0.05; \*\**p* < 0.01; \*\*\**p* < 0.001, \*\*\*\**p* < 0.0001. In I+J, the *p*-values were calculated by two-way ANOVA, followed by pairwise testing with Benjamini-Hochberg correction.

Similarly, we found that *CLU* is expressed at significantly lower levels in LBT003 compared to the other GSCCs (Fig 8A + S8A). As previously mentioned, CLU can act as a ligand of the TREM2 receptor on the surface of GAMs, causing polarization towards an immunosuppressive phenotype. Interestingly, we found that *TREM2* expression in macrophages from the LBT003 co-culture does not significantly differ from the expression in the macrophage monoculture (*p* = 0.8611), while *TREM2* was significantly upregulated in macrophages from the CME037 (*p* = 0.0122), LBT123 (*p* = 0.0028) and CME038 (*p* = 2.8e-8) co-cultures (Fig S8B). The latter two GSCCs were associated with the strongest macrophage polarization towards an immunosuppressive phenotype (see Fig 4D), suggesting that TREM2 signaling may be involved in the polarization process, in addition to *LGALS1*.

To confirm our findings about the top-hit *LGALS1* at the protein level, we evaluated GAL1 expression in the formalin-fixed paraffin-embedded (FFPE) GBM resection specimens from which the GSCCs were derived. Tumor samples from corresponding patients were collected and IHC was used to evaluate the quantity and localization of the GBM tumor cell marker SOX2 and GAL1 (Fig 8C + Supplementary Table 1). We found that GAL1 expression in GBM tumor cells was the lowest for LBT003 compared to GBM tumor tissue from other patients (Fig 8C-D).

Next, we knocked out *LGALS1* in LBT070, the GSCC with the highest GAL1 protein level, using CRISPR-Cas9 technology, and used LBT070 *LGALS1* KO cells in our zebrafish avatar model (Fig 8E + 5I + S8C + Movie 18-19). Strikingly, although the reduction in tumor volume by 5 dpi was comparable for LBT070 and LBT070 *LGALS1* KO (both ±75% reduction) (Fig 8F), LBT070 *LGALS1* KO tumors were much more confined at 5 dpi than the invasive LBT070 tumors (Fig S8D-E). Furthermore, LBT070 *LGALS1* KO tumors attracted significantly more round (*p* = 0.0053), but not ramified macrophages (*p* = 0.6851) at 1 dpi compared to LBT070 tumors (Fig 8G-H). Finally, round macrophages were located significantly closer to LBT070 *LGALS1* KO tumors at 1 dpi (*p* = 1.7e-4), and remained close at 5 dpi, while they moved away from LBT070 tumors (*p* = 1.3e-8) (Fig 8I), suggesting that *LGALS1* KO prevented the switch to a more immunosuppressive phenotype in the GAMs. We observed a similar trend for all macrophages (Fig S8F), but not for the ramified macrophages (Fig 8J). Taken together, these results provide direct evidence that *LGALS1* regulates immunosuppression, and this might be correlated with reduced survival in GBM patients.

## Discussion

Novel therapies for GBM are desperately needed to improve prognosis for this fatal disease. GAMs represent a significant proportion (30-50%) of the tumor and often contribute to immunosuppression and tumor progression, making it a valid target for new therapeutic approaches. In addition to treatments targeting tumor cells, novel immunotherapies have been evaluated and several clinical trials on immune checkpoint inhibitors are ongoing. Furthermore, therapeutic approaches for targeting GAMs, including blocking GAM recruitment, GAM reprogramming and facilitating GAM-mediated phagocytosis are of particular interest. However, these strategies have shown limited efficacy in the clinic so far (Butowski *et al*, 2016). The results obtained in this study, combining scRNA-seq, zebrafish avatars, and IHC, provide a better understanding of the phenotypic alterations and cellular, molecular, and spatial heterogeneity of GSCCs and associated macrophages. Our findings highlight the need for more precise and context-specific interventions that selectively target detrimental immunosuppressive GAMs, while sparing beneficial inflammatory cells.

Using scRNA-seq, we extensively characterized a set of eight diverging GSCCs in an *in vitro* co-culture model with human monocyte-derived macrophages. We reported the molecular characteristics of GAMs and detected patient-specific cell-cell interaction patterns. Even though our findings are based on only eight GSCCs, we uncovered GAM heterogeneity at the molecular level and described a clear phenotype switch, both *in vitro* and *in vivo*. Currently, it is generally accepted that the M1/M2 macrophage terminology originally introduced by Mills and colleagues undermines the complexity of the highly dynamic *in vivo* situation (Mills *et al*, 2000). While the M1/M2 paradigm has been convenient, it has outlived its usefulness in characterizing macrophage subtypes and adds more ambiguity than clarity. With the emergence of single-cell technologies, various studies identified distinct GAM populations and showed that these highly plastic immune cells express a mix of M1 and M2 markers (Abdelfattah *et al*, 2022; Chen *et al*, 2021a; Cui *et al*, 2021; Hara *et al*, 2021; Karimi *et al*, 2023). Concurrent with these observations, we show that macrophages generally polarize towards a more immunosuppressive phenotype upon co-culture with GSCCs, though this shift does not clearly follow the M1/M2 classification. Furthermore, we demonstrate that the degree of macrophage polarization in the co-culture models correlates with GAM recruitment and activity in the corresponding zebrafish avatar models.

We also described the application of advanced live-imaging techniques in an orthotopic zebrafish xenograft model to continuously monitor the dynamic interplay between transplanted GSCCs and GAMs in real-time. In the search for new treatments for GBM, several preclinical models have been developed, including multiple mouse, cell, and organoid models (Mathivet *et al*, 2017; Jacob *et al*, 2020; Zhu *et al*, 2009; Ogawa *et al*, 2018). However, these models do not always accurately reflect the heterogeneity present in the patients, and specifically the complexity of the TME. The use of zebrafish models in cancer research is becoming increasingly popular, due to their unique features like rapid development, small size, transparency of the embryos, ease of genetic manipulation, and ethical and economic advantages (Chen *et al*, 2021b). They have proven highly suited for xenotransplantation of human tumor cells, and GBM tumor cells in particular, due to high similarity with the human brain structure (Reimunde *et al*, 2021). Zebrafish xenograft models are used extensively to study GBM initiation, progression, migration, vasculature and invasion (Pudelko *et al*, 2018; Pan *et al*, 2020; Vittori *et al*, 2016; Umans *et al*, 2021; Mazzolini *et al*, 2022). Hamilton *et al*. studied the interaction between microglia and transplanted GBM cell lines U87 and U251. Concurrent with our results, they observed different growth patterns and microglial responses for both cell lines and confirmed that microglia play a prominent role in promoting GBM tumor cell growth (Hamilton *et al*, 2016). Using an advanced image processing pipeline that was developed in-house, we were able to capture the complex spatiotemporal interaction between GSCCs and associated macrophages. Time-lapse movies uncovered differential tumor cell invasion and infiltration of reactive macrophages across the different avatar models. Using GSEA, we also found that non-invasive GSCCs in the zebrafish avatar model showed an enrichment for ECM production and deposition. This contrasts with the findings of Hoogstrate *et al*., which indicated that an ECM gene signature was associated with significantly worse survival at recurrence, although mainly expressed by pericytes (Hoogstrate *et al*, 2023). The enrichment for OXPHOS in invasive GSCCs, on the other hand, is more evident as motile GBM cells have higher energetic needs (Saurty-Seerunghen *et al*, 2022).

Further, we showed that our models faithfully recapitulated the heterogeneity and clinical features of GBM patients. The rapidly growing field of personalized medicine relies heavily on the use of animal models. Zebrafish avatars of human tumors present a much faster model than murine models and thus hold enormous potential to offer personalized treatment in a clinically relevant time frame. In fact, it has already been demonstrated that zebrafish avatars of various cancer types can predict how individual patients respond to a certain treatment. An example is the study conducted by Fior *et al*., which demonstrated the feasibility of using zebrafish avatars as a tool for testing the efficacy of FOLFOX adjuvant chemotherapy in colorectal cancer patients, as the avatars showed a comparable response to the treatment as their corresponding patients (Fior *et al*, 2017). Although we did not apply therapeutic interventions in this study, our models are ideally suited for pharmaceutical testing and drug discovery. Zebrafish xenograft models have been demonstrated to have clinical features and drug sensitivity similar to human cancers and can thus be used as a fast *in vivo* platform to test existing drugs and to identify relevant novel therapeutic approaches (Usai *et al*, 2020; MacRae & Peterson, 2015; Patton *et al*, 2021). In fact, the zebrafish model has already resulted in the identification of therapeutic targets that were successfully translated to human clinical studies (North *et al*, 2007; Goessling *et al*, 2009; White *et al*, 2011; Owens *et al*, 2008). In the context of GBM, zebrafish xenograft models have become particularly valuable as they allow researchers to assess the ability of drugs to cross the BBB and the presence of systemic toxicity (Zeng *et al*, 2017; Hu *et al*, 2019). Given the substantial heterogeneity and the consistent failure of current therapies in GBM, it is important to explore and develop more individualized GBM treatment strategies. Even though most clinical studies for GBM treatment have failed, there are often exceptional responders who benefit from a particular therapy (Decraene *et al*, 2023). The development and clinical validation of new preclinical models based on patient-derived samples that allow for a more precise reproduction of the patient’s tumor complexity will facilitate a more accurate assessment of whether patients will respond to a specific treatment.

This research identifies Galectin-1 (GAL1, *LGALS1)* as a potential target for immune modulation. We demonstrated that *LGALS1* is involved in immunosuppression, might be correlated with reduced survival in GBM patients, and that *LGALS1* KO in GBM tumor cells can transform the immune landscape. Several research groups have reported similar findings (Chou *et al*, 2018; Chen *et al*, 2019; Sharanek *et al*, 2021). *LGALS1* has been shown to drive resistance to chemo- and immunotherapy, and to be associated with poor survival. Moreover, chitosan nanoparticles loaded with siRNA targeting GAL1 have been developed for intranasal delivery to the tumor and its microenvironment (Van Woensel *et al*, 2016; Van Woensel *et al*, 2017). Treatment with these nanoparticles reduced immunosuppression and increased survival in tumor-bearing mice. The authors also described synergistic effects for the combination of anti-GAL1 treatment with chemo- and immunotherapy, which indicates that anti-GAL1 nanoparticles could be a valuable adjuvant treatment. In summary, these observations strongly suggest that incorporating GAL1-targeted therapeutics into the current treatment schedule of GBM patients could be an effective approach for enhancing the outcome of GBM patients.

Furthermore, this study highlights a potential role for TREM2 signaling in GBM disease progression by creating an immunosuppressive environment. Indeed, by predicting L:R interactions between GBM tumor cells and monocyte-derived macrophages, we revealed that there was more putative TREM2 signaling in co-cultures that exhibited strong macrophage polarization. TREM2 signaling is known to be involved in many different pathologies, including neurodegenerative disease (Li & Zhang, 2018), obesity (Reich *et al*, 2023) and several cancers (Deczkowska *et al*, 2020; Katzenelenbogen *et al*, 2020), where it plays an important role in immunosuppression. In the context of GBM, one study indicated that *TREM2* expression was correlated with poor tumor immunity and worse prognosis, and that knock-down of *TREM2* resulted in a decrease in M2 polarization (Yu *et al*, 2023). Further studies remain necessary to elucidate detailed mechanisms underlying the immunosuppressive role of both *LGALS1* and TREM2 signaling.

Taken together, our scRNA-seq profiling assay complemented with our zebrafish avatar model provide remarkable ability to reveal heterogeneity in GBM tumor growth and interactions within the TME. They can be employed in preclinical research to better mimic GBM and the interactions of GAMs, which could be a crucial step towards improved GBM treatments. Furthermore, we envision our models to act as the foundation for developing a functional screening platform to identify promising immune-modulating targets. Finally, our fast and sensitive models hold enormous potential to maximize therapy efficacy in a personalized fashion. They could be used to predict patient’s response to certain treatments, as well as to establish better inclusion criteria for clinical trials, resulting in higher success rates.

## Materials and Methods

### Patient-derived GBM stem cell cultures

Fresh brain tumor tissues were obtained with informed consent from patients undergoing surgical resection. Samples were processed into single-cell suspensions immediately after resection. The tumor tissue was cut into pieces of 1-2 mm that were mechanically and enzymatically dissociated with the MACS Brain Tumor Dissociation Kit, gentleMACS Dissociator and MACSmix Tube Rotator (Miltenyi Biotec) according to the manufacturer’s protocol. The dissociated sample was passed through a 70 µm strainer to remove remaining larger particles. ACK lysing buffer was added to eliminate the red blood cells. After centrifugation, the cell pellet was resuspended in NeuroCult NS-A Basal Medium (human, STEMCELL Technologies) supplemented with NeuroCult NS-A Proliferation Supplement (human, STEMCELL Technologies), heparin solution (2 mg/mL, STEMCELL Technologies), recombinant bFGF (20 ng/mL, human, STEMCELL Technologies), recombinant EGF (20 ng/mL, human, STEMCELL Technologies) and antibiotic antimycotic solution (100x; 5 ml, Sigma-Aldrich), hereafter referred to as complete GBM medium. Live cells were counted using an automated cell counter (Bio-Rad). Part of the cells were frozen, and the rest was plated in an ultra-low attachment flask (Corning) and/or a cell culture flask (SARSTEDT) coated with laminin (Sigma-Aldrich). The medium was changed twice a week. Cells were split when 80-90% confluency was reached. This study and procedures have been evaluated and approved by the Ethical Committee Research of UZ Leuven / KU Leuven (S59804 & S61081). Generated GSCCs were characterized as previously described (Rosenberg et al, 2017). BT333 and BT569 were obtained from Dana-Farber Cancer Institute (Boston, MA, USA) (IRB 10-417).

### Monocyte isolation

Blood was obtained from healthy donors with informed consent and with approval from the Ethical Committee Research of UZ Leuven / KU Leuven (S59804 & S61081) and collected in cell preparation tubes (CPT – Sodium Heparin, BD Biosciences). From the blood, peripheral blood mononuclear cells (PBMCs) were isolated according to the manufacturer’s protocol (CPT, BD Biosciences). Next, monocytes were isolated from the PBMCs using a monocyte isolation kit (Pan Monocyte Isolation Kit, human, Miltenyi Biotec). Next, 1x10^7^ cells were plated in a T25 cell culture flask and cultured in RPMI 1640 medium with GlutaMAX (Gibco) supplemented with 50 ng/ml GM-CSF (Recombinant human, R&D Systems), 10% heat-inactivated FBS (Gibco), and antibiotic antimycotic solution (100x, 5 ml), hereafter referred to as complete mφ medium. The rest of the monocytes was frozen in aliquots of 10x10^6^ cells. Monocytes were cultured and differentiated into macrophages as previously described (Jin & Kruth, 2016).

### Cell culture

All cells were maintained at 37°C and 5% CO_2_. GBM cells and monocytes/macrophages were cultured in complete GBM medium and complete mφ medium, respectively. LBT001, LBT003, LBT070, LBT123, CME037 and CME038 were grown on laminin-coated flasks. BT333 and BT569 were grown on ultra-low attachment plates. The medium was changed two to three times a week and cells were passaged when 80-90% confluency was reached. For passaging, GBM cells growing on laminin-coated plates were collected using a cell scraper, GBM cells growing on ultra-low attachment plates were collected through centrifugation, and macrophages were harvested using 0.25% trypsin-EDTA (Gibco), after washing with PBS (Gibco) without CaCl_2_ and MgCl_2_. For cell counting, a single-cell suspension was created by treating the GBM cells for 3 minutes with accutase (StemPro Accutase Cell Dissociation Reagent, Gibco) at 37°C.

### Co-culture model of human macrophages and GBM cells

To evaluate the cell-cell interaction between GBM cells and macrophages, co-culture experiments were carried out as follows: 1x10^4^ monocyte-derived macrophages were seeded together with 4x10^4^ GBM cells in hanging drops of 20 μl complete GBM medium (Keller, 1995; Foty, 2011). As a control, macrophage monoculture was established by seeding 2.5 x10^4^ macrophages in hanging drops of 20 μl complete GBM medium. The number of droplets per GSCC ranged from 115 to 139. The cells were co-cultured for four days. At the end of the experiment, the cells were labeled with MULTI-seq for scRNA-seq.

### MULTI-SEQ

MULTI-seq was used to label cells with sample-specific barcodes prior to pooling and single-cell RNA sequencing (McGinnis *et al*, 2019). Cells were prepared as follows. Cells were washed with PBS. Macrophages were treated with 0.25% trypsin-EDTA for 10 minutes at 37°C. Co-cultures were first treated with accutase for 3 minutes at 37°C to collect the GBM cells and subsequently with 0.25% trypsin-EDTA for 15 minutes at 37°C to collect the macrophages. Single-cell suspensions were then pelleted for 5 minutes at 300 g and washed twice with PBS. Cells were pelleted and resuspended in 200 ml of a 200 nM solution containing equimolar amounts of anchor LMO and sample barcode oligonucleotides in PBS and incubated on ice for 5 minutes. Next, 20 μL of a 2 μM co-anchor LMO solution in PBS (for a final concentration of 200 nM) was added to each sample. Following gentle mixing, the labeling reaction was continued on ice for another 5 minutes. For the remaining procedure, cells were kept on ice. Then, 1 ml of ice cold 1% BSA in PBS was added and cells were pelleted at 4°C. Next, labeled cells were washed twice with ice cold 1% BSA in PBS, counted and pooled into a single aliquot. Pooled sample was concentrated to 10^6^ cells/ml in 1% BSA in PBS. Subsequently, the scRNA-seq procedure was followed.

### Single-cell RNA sequencing & analysis

The pooled cell suspension containing several samples individually barcoded by MULTI-seq was processed with 10x Genomics Technology. The link between the 10x cell barcode and MULTI-seq barcode was achieved using the deMULTIplex R package (https://github.com/chris-mcginnis-ucsf/MULTI-seq). Cells without viable MULTI-seq barcode or with ambiguous MULTI-seq barcode were discarded. Cell-barcoded 5′ gene expression libraries were sequenced on an Illumina NovaSeq6000 system and mapped to the GRCh38 human reference genome using CellRanger (10x Genomics). Raw gene expression matrices were generated using CellRanger. Downstream analysis was performed with the Seurat R package (Satija *et al*, 2015; Butler *et al*, 2018; Stuart *et al*, 2019; Hao *et al*, 2021). Cells with unambiguous MULTI-seq barcode, expressing >200 and <6000 genes, containing >400 unique molecular identifiers (UMIs) and <20% mitochondrial counts were retained in the analysis. After normalization and regression for the number of UMIs, percentage of mitochondrial genes and cell cycle (S and G2M scores were calculated by the *CellCycleScoring* function in Seurat), the PCs were calculated based on the 2000 most variable genes. The PCs explaining the highest variance were used for clustering (SNN + Louvain) (Blondel *et al*, 2008; Ertöz *et al*, 2003) and visualization with UMAP (McInnes *et al*, 2018). Clusters in the resulting two-dimensional UMAP representation consisted of distinct cell types, which were identified based on the expression of marker genes. Differential expressed genes were identified using the MAST test (Finak *et al*, 2015) with *FindMarkers* and *FindAllMarkers* functions in Seurat. Following manual annotation of GBM cells and macrophages, based on expression of *SOX2* and *CD68* respectively, subclustering of macrophages only and GBM cells only was performed using the same procedures as described above (2000 most variable genes, selection of most relevant PCs, UMAP).

Following SNN clustering and Louvain community detection, 3 distinct clusters were identified in the macrophage dataset. Following marker analysis using the *FindAllMarkers* functions in Seurat, one cluster was identified as transitioning monocytes (MC3). For subsequent pseudotime analyses, this cluster was removed, and subclustering was performed again with the remaining two clusters, following the same steps described above. PCA revealed that the bulk of the variance between the two remaining clusters (MC1/MC2) was defined by PC1.

As such, the PC1 value was used as a measure of pseudotime along the MC1-MC2 axis. To identify temporally expressed genes, a general additive model (GAM) was fitted for the 1000 most variable genes with a locally estimated scatterplot smoothing (LOESS) term for the PC1 value, using the tradeSeq R package (*fitGAM* function, n = 4 knots) (Van den Berge *et al*, 2020). A heatmap of the 100 most significantly temporally expressed genes (as identified with the *associationTest* function) was generated using the *predictSmooth* function (nPoints = 50) and the pheatmap R package (https://CRAN.R-project.org/package=pheatmap).

CellPhoneDB analysis was performed to detect putative L:R interactions between GSCCs and macrophages (and vice versa) (Vento-Tormo *et al*, 2018; Efremova *et al*, 2020). CellPhoneDB package v4.0.0 (Garcia-Alonso *et al*, 2022) was used to perform pairwise comparison between GBM cells and macrophages (and vice versa) by randomly permuting labels of the clusters 1000 times (default setting), thus generating a null-distribution of each L:R pair, and then determining the actual mean log2 expression of the L:R pair. Of note, only receptors and ligands that were detected in at least 10% of the cells in each respective cluster of interest were considered for the analysis (default setting). *P*-values for each interaction were then determined by comparing the proportion of the means that were equal to or higher than the actual mean of a given L:R pair. Only interactions with a *p*-value ≦ 0.05 were included in the figures.

Subclustering of GBM cells was performed as described above (regression of cell cycle, selection of 2000 most variable genes, selection of most relevant PCs, UMAP). Distinct markers for LBT003 were determined using the MAST test (Finak *et al*, 2015) with the *FindMarkers* function in Seurat. Stratification of the single GBM cells into the Neftel subtypes was performed as described by Neftel *et al*. (Neftel *et al*, 2019).

GSEA was performed with the *clusterProfiler* (version 3.16) package in R. Canonical pathway gene sets were downloaded from MSigDB (Subramanian *et al*, 2005), which combines gene sets from the WikiPathways, Reactome, KEGG, PID and BioCarta databases. Gene sets with a size <15 or >500 genes were excluded from the analysis. A ranked list of log2(FC) was used as an input of the *GSEA* function of the *clusterProfiler* package. The 25 most significant pathways (ranked by corrected *p*-value) were subsequently visualized with the *dotplot* function.

### Transduction (GFP labeling & *LGALS1* knock-out)

For visualization in the zebrafish avatar model, GBM cells were stably labeled with GFP using lentiviral transduction (PD-18-046: pCH-EF1a-eGFP-IRES-Puro, 1.18 x 10^8^ TU/ml, Leuven Viral Vector Core (LVVC)). For *LGALS1* KO in GBM cells, CRISPR/Cas9 technology was used. All-in-one plasmid was adapted from pXPR023 (Broad Institute) by replacing the puromycin resistance gene with eGFP. Expression of humanized *S. pyogenes* Cas9 was driven by the EFS-NS promoter in a bicistronic cassette using eGFP as a reporter gene. Four guide RNAs (gRNAs) were designed using the online tool of Broad Institute (https://portals.broadinstitute.org/gpp/public/analysis-tools/sgrna-design) to maximize the chances of effective *LGALS1* KO (Supplementary Table 1). Single gRNA sequences were integrated in the plasmid under control of the human U6 promoter, yielding four different plasmids. Lentiviral particles were generated from an equimolar mixture of the plasmids by LVVC (PD-22-119: pXPR023-eGFP, 1.03 x 10^7^ TU/ml). Cells with most optimal GFP labeling were grown and sorted using FACS to exclude non-transduced cells. Cells were sorted with a Sony MA900 cell sorter from the KU Leuven Flow and Mass Cytometry Facility.

### Animal care and handling

*In vivo* experiments were conducted in zebrafish (*Danio rerio*). Adult fish were maintained under standard laboratory conditions (Nüsslein-Volhard & Dahm, 2002). Zebrafish embryos were raised and staged as previously described (Kimmel *et al*, 1995). All zebrafish were maintained and handled in compliance with European Animal Welfare Legislation (2010/63/EU) and FELASA guidelines and recommendations concerning laboratory animal welfare and scientific use. All protocols involving work with live animals that are described below were reviewed and approved by the Animal Ethics Committee of the KU Leuven (P167/2021). The experiments of this project were performed by LF, who holds a FELASA B certificate, in the zebrafish facility and laboratory at VIB – KU Leuven.

### Transgenic and mutant zebrafish lines

*Tg(mpeg1:mCherryF)^ump2^* zebrafish embryos were used for the visualization of macrophages (and microglia) (Ellett *et al*, 2011; Silva *et al*, 2021). The vascular reporter line *Tg(kdrl:lynEYFP)* was generated in the mutant transparent *casper* background for improved *in vivo* visualization (White *et al*, 2008). Growing and breeding of transgenic and mutant lines was done in accordance with the regulations of the Animal Ethics Committee of the KU Leuven.

### Zebrafish transgenesis

The construct for transgenesis of the *Tg(kdrl:lynEYFP)* line was generated by cutting and ligating the *kdrl* promoter and the coding sequence of lynEYFP, making use of adjacent restriction sites present in the vectors. The final construct was ligated into a Tol2 destination vector (pDestTol2pACryGFP, Supplementary Table 1) (Kwan *et al*, 2007). Transgenesis was done via Tol2-mediated recombination as previously described (Kwan *et al*, 2007; Kawakami, 2004). *Casper* embryos were co-injected at one-cell stage with 100 pg of Tol2 mRNA and 40 pg of pDestTol2-kdrl-lynEYFP plasmid DNA. Embryos were raised at 28°C and screened for EYFP expression at ∼72 hpf.

### Orthotopic zebrafish xenograft model

Heterozygous mutant *casper* embryos were treated with 0.003% 1-phenyl-2-thiourea (PTU, Sigma-Aldrich) from 24 hpf onwards to delay pigmentation. At 30 hpf, GFP-expressing GBM tumor cells were microinjected into the hindbrain ventricle of anesthetized (0.014% tricaine, MS-222, Sigma-Aldrich) *Tg(mpeg1:mCherryF)^ump2^; Tg(kdrl:lynEYFP)* embryos. Approximately 300-800 cells were injected per embryo and zebrafish avatars were grown at 34°C until the end of the experiment. At 1 dpi (54 hpf), zebrafish avatars were screened for the presence of GBM cells and those with the largest tumors were selected for overnight imaging. At 5 dpi (144 hpf), the same avatars were imaged again to follow tumor progression.

### Live-imaging

Embryos were anesthetized in 0.014% tricaine, mounted in a 35 mm glass bottom petri dish (0.17 mm, MatTek) using 0.6% low-melting-point agarose (Lonza) containing 0.014% tricaine, and bathed in Danieau buffer containing 0.007% tricaine and 0.003% PTU. Time-lapse imaging was performed using a Leica TCS SP8 upright microscope with a Leica HCX IRAPO L x25/0.95 water-dipping objective and heating chamber. Zebrafish avatars were typically imaged at 34°C at two different timepoints: 1 dpi and 5 dpi. Time-lapse images were acquired for up to 16 hours at 1 dpi and up to 8 hours at 5 dpi and time intervals ranged from 15 to 30 minutes.

### Image processing

The 3D time-lapse movies were processed by image analysis and machine learning algorithms to faithfully segment tumor and macrophages over time and classify GSCC-specific morphometrics and dynamics of the tumor and its microenvironment. Our dataset contains 190 3D time-lapse movies consisting of three channels: GBM tumor cells, macrophages, and vasculature. Image pre-processing was performed using Fiji (version 2.0.0) (Schindelin *et al*, 2012), including conversion to tiff image file format and separation of the channels. Bleed-through of the tumor signal into the vascular channel was removed. Autofluorescence correction proved necessary especially for 5 dpi movies due to increased pigmentation of the zebrafish embryos and was achieved using ilastik’s pixel classification algorithm (Berg *et al*, 2019), which was trained on annotations of tumor, macrophages, autofluorescence and background. After obtaining the classification, the pixels containing autofluorescence were extracted and removed. Following autofluorescence removal, the tumor and macrophage channels were analyzed using a customized image analysis pipeline. The tumor channel was denoised using a Gaussian blur filter and segmented by Otsu thresholding (Otsu, 1979). For macrophage segmentation, the macrophage channel was initially max-intensity projected in 2D. The 2D macrophage images were segmented using the deep learning algorithm Cellpose (Pachitariu & Stringer, 2022). The Cellpose model was trained using more than 1000 manually annotated macrophages on 42 2D images, equally distributed over movies starting at 1 dpi and 5 dpi, and all GSCCs. Subsequently, the Cellpose model was applied to all movies to obtain a segmentation of the single macrophages. Macrophage segmentation was validated in a subset of movies by comparing Cellpose-segmented macrophage masks with manually annotated macrophage masks. The quality of the segmentation was assessed by identifying correctly segmented macrophages versus false positives and false negatives, based on optimizing the intersection over union (IoU) metric. Subsequently, a 3D from 2D inference for the center positions of the macrophages was applied to identify 3D positions based on the macrophage segmentations in 2D. This workflow is an Album solution (Albrecht *et al*, 2021) accessible on <u>“Album solution”</u>. To compute distances of the macrophages to the tumor, a Euclidean distance transform was applied on the 3D tumor masks. This allowed us to obtain the shortest distance of each macrophage center position to the surface of the tumor. For macrophages in contact with or inside the tumor the distance is 0. Finally, automated quantitative analysis was performed whereby various metrics were obtained from the segmentation masks (tumor volume, macrophage number, macrophage shape, *etc.*) by using scikit-image, a collection of image processing algorithms in Python (van der Walt *et al*, 2014). Macrophage circularity was computed as follows: 4*πA*/*p*^2^, where *A* is the area of the macrophage and *p* its perimeter. A circularity of 1 indicates a perfect circle, while circularity values close to 0 indicate highly non-circular shapes. Round macrophages were defined as macrophages with a circularity larger than 0.6, while ramified macrophages were defined as cells with a circularity smaller than 0.35. The analysis scripts implemented in Python are available in a <u>GitHub repository</u>. For plotting time courses of different features, an R Shiny app was used, which was derived from PlotTwist (Goedhart, 2020) and can be found in the same repository.

### Immunohistochemistry

FFPE GBM tissue samples (collected at the UZ/KU Leuven biobank according to protocols S59804 and S61081) were used for IHC. Patient material was available for all patients, except for BT333 and BT569. If sufficient material was available, multiple regions were selected per tumor sample, and a tissue microarray (TMA) was created. Our TMA included 15 cores in total, originating from four patients (LBT001, LBT003, CME037 and CME038). For LBT070 and LBT123, whole-slide (WS) sections were used due to limited tissue availability.

First, FFPE tissue slides were deparaffinized by sequentially placing them in xylene, 100% ethanol and 70% ethanol. Following dewaxing, antigen retrieval was performed with PT link (Agilent) using 10 mM EDTA in Tris-buffer (pH 8). Next, slides were placed in bleaching buffer to lower the intrinsic autofluorescent background signal. Immunofluorescence staining was performed using Bond RX Fully Automated Research Stainer (Leica Biosystems) with anti-GAL1 (Cell Signaling Technology, 13888, 1:375) and anti-SOX2 (ThermoFisher Scientific, 14-9811-82, 1:150) antibodies (Supplementary Table 1). Tissue sections were incubated for 4 hours with the previously validated primary antibodies, washed, and then incubated with fluorescently labeled secondary antibodies for 30 minutes. A coverslip was placed onto the slides with medium containing DAPI to stain cell nuclei. A high-resolution image was generated at ×10 magnification using a Axio Scan.Z1 slide scanner (Zeiss).

Raw scans were transformed to grayscale 16-bit tiff images using the developer’s software (Zen). Image registration was performed by applying a homomorphic transformation over a set of matched descriptors using a Harris detector. Images were subsequently adjusted for background intensity variations using the rolling ball algorithm, and autofluorescence removal was performed by subtracting the pre-stained image of the corresponding tissue section from the measured signal. Intensities for both images were normalized using quantile normalization. Mean fluorescence intensity values for GAL1 staining in SOX2^+^ cells were normalized using *Z*-scores within each core/WS and averaged per patient, as previously described (Caicedo *et al*, 2017). To avoid a strong influence from outliers, *Z*-scores were trimmed within the [-5, 5] range. For enhanced visualization, a binary mask was generated of the SOX2^+^ cells and multiplied with the image of GAL1 staining in Fiji (version 2.0.0) to exclude GAL1 staining in non-tumor cells (see Fig 8C).

LBT070 cells and LBT070 LGALS1 KO cells were stained with anti-GAL1 antibody (Cell Signaling Technology, 13888, 1:375) and staining was evaluated using the Operetta High Content Imaging System (PerkinElmer) (See Fig S8C).

### Statistics

Statistical details of all analyses are reported in the figure legends. All statistical tests used were two-sided unless otherwise mentioned. All reported measurements were taken from distinct samples and not repeatedly measured on the same sample, unless at different timepoints (1 dpi and 5 dpi). Tumor volumes at 1 dpi and 5 dpi (Fig S6B-C) were normally distributed as confirmed by Shapiro-Wilk normality tests. According to the central limit theorem, we assumed that macrophage distances were normally distributed (*n* > 30). For all other statistical analyses of *in vivo* data, two-way repeated measures ANOVA (or mixed-effects model in case of missing values) was used, for which no non-parametric equivalent exists. All statistical analyses or graphical representations were executed using R Studio version 2022.12.0+353, or GraphPad Prism version 9.5.1.

## Data Availability

Image analysis scripts are available on GitHub: https://github.com/wgiese/zebrafish_xenograft_analysis

## Acknowledgements

This work was supported by FWO (Fonds voor Wetenschappelijk Onderzoek – Vlaanderen / Flemish Fund for Scientific Research) grants (G0B3722N, G0I1118N and S001221N; FDS, BC, MD, GS), a KOTK (Kom op tegen Kanker / Stand against Cancer) grant (KOTK/2019/11892/1; FDS), KU Leuven grants (IDN/20/021 and CELSA/20/022; FDS), Foundation Leducq Transatlantic Network of Excellence Grant ATTRACT (17 CVD 03; HG), and Deutsche Forschungsgemeinschaft (DFG) (GE2154/1-1; HG, EB). LF was supported by an FWO PhD fellowship strategic basic research (1S49718N & 1S49720N). MV is supported by an FWO PhD fellowship fundamental research (11L0822N). BD and SDV are supported by a KOTK grant (KOTK/2018/11509/2). MNK is supported by iNAMES (MDC-Weizmann Research School, Imaging from NAno to MESo). JPA is funded by Helmholtz Imaging, a platform of the Helmholtz Incubator on Information and Data Science. KLL acknowledges support from NIH P50CA165962, R01CA188228, R01CA262462, R01CA215489, and R01CA219943. BB and DL are funded by FWO and the Flemish Government.

We thank Christopher S. McGinnis for his advice on MULTI-seq methodology and demultiplexing. *Tg(mpeg1:mCherryF)^ump2^*zebrafish embryos were kindly provided by Prof. Jean-Pierre Levraud. Lentiviral particles (GFP labeling and *LGALS1* KO) were generated by Leuven Viral Vector Core. Figure 1A, 2A, 5A and 6A were created with BioRender.com.

## Author contributions

**Lise Finotto:** Conceptualization; data curation; formal analysis; funding acquisition; investigation; methodology; project administration; validation; visualization; writing – original draft; writing – review and editing. **Basiel Cole:** Data curation; formal analysis; validation; visualization; writing – review and editing. **Wolfgang Giese:** Data curation; formal analysis; methodology; software; validation. **Elisabeth Baumann:** Data curation; formal analysis; methodology; software; validation. **Annelies Claeys:** Investigation; validation. **Maxime Vanmechelen:** Investigation. **Brecht Decraene:** Investigation. **Marleen Derweduwe:** Investigation. **Nikolina Dubroja Lakic:** Investigation. **Gautam Shankar:** Formal analysis; software. **Madhu Nagathihalli Kantharaju:** Software. **Jan Philipp Albrecht:** Software. **Ilse Geudens:** Conceptualization; investigation. **Fabio Stanchi:** Investigation. **Keith L. Ligon:** Resources. **Bram Boeckx:** Resources. **Diether Lambrechts:** Resources. **Kyle Harrington:** Methodology; software. **Ludo Van Den Bosch:** Resources. **Steven De Vleeschouwer:** Resources. **Frederik De Smet:** Conceptualization; funding acquisition; methodology; project administration; supervision; writing – review and editing. **Holger Gerhardt:** Conceptualization; funding acquisition; methodology; project administration; supervision; writing – review and editing.

## Table legends

**Supplementary Table 1: Overview of gRNAs, plasmids and antibodies**

A gRNA sequences used for *LGALS1* KO.

B Plasmids used to create viral particles for *LGALS1* KO and for generating a vascular reporter line in zebrafish.

C Antibodies and associated details used for IHC.

## Expanded View Figure legends

**Supplementary Figure 2: Single-cell profiling of G*SCC*-macrophage co-cultures.**

A Violin plot showing the number of detected genes per cell. (*n* = 5320 cells from 9 samples with a median of 3334 genes detected per cell.)

B-C UMAP plots showing *SOX2* (B) and *CD68* (C) expression.

D UMAP plot showing cell cycle score. Proliferating GBM tumor cells are depicted in blue.

**Supplementary Figure 3: Molecular heterogeneity of GBM-associated macrophages.**

A UMAP plot of macrophage population identified three distinct macrophage subclusters (MC1-3).

B Macrophage subcluster distribution for the different samples.

**Supplementary Figure 4: Macrophages shift towards an immunosuppressive phenotype upon co-culture with patient-derived GSCCs.**

A UMAP plot of macrophage population without TransMos shows two distinct macrophage subclusters (MC1-2).

B Representation of original samples on UMAP plot.

C-G LOESS plots for *AKR1B1* (C), *CCL4* (D), *MSR1* (E), *LIPA* (F), and *LGALS1* (G).

H Dot plot of cell–cell communication analysis using CellPhoneDB. Depicted are L:R pairs for macrophage – GSCC signaling across all GSCCs, ranked by mean log2 expression. Each dot size shows the log2 mean of expression values and dot color indicates the *p*-value for the listed L:R pairs (x-axis) in the respective GSCCs (y-axis). Only top 50 significant L:R pairs, with cut-offs of *p*-value ≦ 0.05 are shown. The *p*-values were generated by CellPhoneDB, which uses a one-sided permutation test to compute significant interactions.

**Supplementary Figure 5: Overview of different zebrafish avatars.**

A Number of zebrafish embryos used in the time-lapse movies that were generated at 1 dpi and 5 dpi.

B Representative maximum-projections of a z stack of the head region of a *Tg(mpeg1:mCherryF)^ump2^; Tg(kdrl:lynEYFP)* zebrafish embryo with a GFP-labeled LBT070 tumor, at 37.5, 38, 38.5 and 39 hpi to illustrate the phagocytosis of a GBM tumor cell by a round macrophage (indicated by arrows). GBM tumor cells are shown in green, macrophages in red, and blood vessels in blue. Scale bars: 50 μm.

**Supplementary Figure 6: GSCC-specific morphometrics and dynamics of the tumor and its microenvironment in 3D over time.**

A Mean tumor volume over time, during 1 dpi movies (left) and 5 dpi movies (right). (*n* = 4-22 zebrafish embryos per group, with an average of 10.81 embryos per group).

B-C Tumor volume at the start of 1 dpi (B) and 5 dpi (C) time-lapse movies. (*n* = 4-22 zebrafish embryos per group, with an average of 10.81 embryos per group).

D-E Trend line of median macrophage distance to the tumor at 1 dpi (D) and 5 dpi (E) for all GSCCs.

F Boxplot of distance of round and ramified macrophages to the tumor for all GSCCs, at the start of 1 dpi and 5 dpi time-lapse movies. (*n* = 491-2238 macrophages per group, with an average of 1330 macrophages per group; boxes stand for 50% of the data and minima/maxima are indicated by the line ends).

G Boxplot of macrophage distance to the tumor of round macrophages within 30 μm of the tumor, at the start of 1 dpi and 5 dpi time-lapse movies, ranked by increasing median distance at 1 dpi. (*n* = 11-324 macrophages per group, with an average of 103.44 macrophages per group; boxes stand for 50% of the data and minima/maxima are indicated by the line ends).

H Boxplot of macrophage distance to the tumor of ramified macrophages within 30 μm of the tumor, at the start of 1 dpi and 5 dpi time-lapse movies, ranked by increasing median distance at 1 dpi. (*n* = 3-86 macrophages per group, with an average of 29.31 macrophages per group; boxes stand for 50% of the data and minima/maxima are indicated by the line ends).

Data information: In B-C, data are presented as mean ± SD. The *p*-values were calculated by one-way ANOVA, followed by Tukey’s multiple comparisons correction. ns ≥ 0.05, \**p* < 0.05; \*\**p* < 0.01; \*\*\**p* < 0.001, \*\*\*\**p* < 0.0001. In D+E, the *p*-values were calculated by the Mann Kendall test. In F, the *p*-values were calculated by two-way ANOVA.

**Supplementary Figure 7: Macrophage/GAM-GSCC interactions correlate to clinical outcome in GBM patients.**

A UMAP plot of GBM tumor cells showing cell cycle score.

B UMAP plot of GBM tumor cells showing Neftel subtypes.

**Supplementary Figure 8: *LGALS1* is involved in suppression of the immune system.**

A-B Violin plots showing *CLU* (A) and *TREM2* (B) expression levels in GSCCs (A) and macrophages (B). C Representative immunofluorescence images of LBT070 *LGALS1* WT and LBT070 *LGALS1* KO cells showing expression of GAL1 (green), and cell nuclei stained by DAPI. Scale bars: 100 μm.

D-E Representative maximum-projections of a z stack of the head region of *Tg(mpeg1:mCherryF)^ump2^; Tg(kdrl:lynEYFP)* zebrafish embryos with GFP-labeled LBT070 (D) and LBT070 *LGALS1* KO (E) tumors at 5 dpi. GBM tumor cells are shown in green, macrophages in red, and blood vessels in blue. Scale bars: 50 μm. *n* = 9 (D) and 5 (E).

F Boxplot of macrophage distance to the tumor of all macrophages within 30 μm of the tumor, at the start of 1 dpi and 5 dpi time-lapse movies. (*n* = 83-481 macrophages per group, with an average of 230 macrophages per group; boxes stand for 50% of the data and minima/maxima are indicated by the line ends).

Data information: In B, the *p*-values were calculated by Kruskal-Wallis test (*p* = 2.5e-10). ns ≥ 0.05, \**p* < 0.05; \*\**p* < 0.01; \*\*\**p*

< 0.001, \*\*\*\**p* < 0.0001. In F, the *p*-values were calculated by two-way ANOVA, followed by pairwise testing with Benjamini-Hochberg correction.

## Movie legends

**Movie 1-2:** Representative time-lapse movies of maximum-projected z stacks of the head region of a *Tg(mpeg1:mCherryF)^ump2^; Tg(kdrl:lynEYFP)* zebrafish embryo with a GFP-labeled BT333 tumor at 1 dpi (Movie 1) and 5 dpi (Movie 2). GBM tumor cells are shown in green, macrophages in red, and blood vessels in yellow. Scale bar = 50 μm, time is indicated in hours:minutes, time interval (T_i_) = 15 min, frame rate = 4 frames per second (fps).

**Movie 3-4:** Representative time-lapse movies of maximum-projected z stacks of the head region of a *Tg(mpeg1:mCherryF)^ump2^; Tg(kdrl:lynEYFP)* zebrafish embryo with a GFP-labeled BT569 tumor at 1 dpi (Movie 3) and 5 dpi (Movie 4). GBM tumor cells are shown in green, macrophages in red, and blood vessels in yellow. Scale bar = 50 μm, time is indicated in hours:minutes, T_i_

= 20 min, frame rate = 4 fps.

**Movie 5-6:** Representative time-lapse movies of maximum-projected z stacks of the head region of a *Tg(mpeg1:mCherryF)^ump2^; Tg(kdrl:lynEYFP)* zebrafish embryo with a GFP-labeled CME037 tumor at 1 dpi (Movie 5) and 5 dpi (Movie 6). GBM tumor cells are shown in green, macrophages in red, and blood vessels in yellow. Scale bar = 50 μm, time is indicated in hours:minutes, T_i_ = 20 min, frame rate = 4 fps.

**Movie 7-8:** Representative time-lapse movies of maximum-projected z stacks of the head region of a *Tg(mpeg1:mCherryF)^ump2^; Tg(kdrl:lynEYFP)* zebrafish embryo with a GFP-labeled CME038 tumor at 1 dpi (Movie 7) and 5 dpi (Movie 8). GBM tumor cells are shown in green, macrophages in red, and blood vessels in yellow. Scale bar = 50 μm, time is indicated in hours:minutes, T_i_ = 20 min, frame rate = 4 fps.

**Movie 9-10:** Representative time-lapse movies of maximum-projected z stacks of the head region of a *Tg(mpeg1:mCherryF)^ump2^; Tg(kdrl:lynEYFP)* zebrafish embryo with a GFP-labeled LBT001 tumor at 1 dpi (Movie 9) and 5 dpi (Movie 10). GBM tumor cells are shown in green, macrophages in red, and blood vessels in yellow. Scale bar = 50 μm, time is indicated in hours:minutes, T_i_ = 20 min, frame rate = 4 fps.

**Movie 11-12:** Representative time-lapse movies of maximum-projected z stacks of the head region of a *Tg(mpeg1:mCherryF)^ump2^; Tg(kdrl:lynEYFP)* zebrafish embryo with a GFP-labeled LBT003 tumor at 1 dpi (Movie 11) and 5 dpi (Movie 12). GBM tumor cells are shown in green, macrophages in red, and blood vessels in yellow. Scale bar = 50 μm, time is indicated in hours:minutes, T_i_ = 20 min, frame rate = 4 fps.

**Movie 13-14:** Representative time-lapse movies of maximum-projected z stacks of the head region of a *Tg(mpeg1:mCherryF)^ump2^; Tg(kdrl:lynEYFP)* zebrafish embryo with a GFP-labeled LBT070 tumor at 1 dpi (Movie 13) and 5 dpi (Movie 14). GBM tumor cells are shown in green, macrophages in red, and blood vessels in yellow. Scale bar = 50 μm, time is indicated in hours:minutes, T_i_ = 30 min, frame rate = 4 fps.

**Movie 15-16:** Representative time-lapse movies of maximum-projected z stacks of the head region of a *Tg(mpeg1:mCherryF)^ump2^; Tg(kdrl:lynEYFP)* zebrafish embryo with a GFP-labeled LBT123 tumor at 1 dpi (Movie 15) and 5 dpi (Movie 16). GBM tumor cells are shown in green, macrophages in red, and blood vessels in yellow. Scale bar = 50 μm, time is indicated in hours:minutes, T_i_ = 20 min, frame rate = 4 fps.

**Movie 17:** Representative time-lapse movie of maximum-projected z stacks of the head region of a *Tg(mpeg1:mCherryF)^ump2^; Tg(kdrl:lynEYFP)* zebrafish embryo with a GFP-labeled LBT070 tumor at 1 dpi to illustrate the phagocytosis of a GBM tumor cell by a round macrophage (indicated by arrows from 12:00 – 15:00). GBM tumor cells are shown in green, macrophages in red, and blood vessels in yellow. Scale bar = 50 μm, time is indicated in hours:minutes, T_i_ = 20 min, frame rate = 4 fps.

**Movie 18-19:** Representative time-lapse movies of maximum-projected z stacks of the head region of a *Tg(mpeg1:mCherryF)^ump2^; Tg(kdrl:lynEYFP)* zebrafish embryo with a GFP-labeled LBT1070 *LGALS1* KO tumor at 1 dpi (Movie 18) and 5 dpi (Movie 19). GBM tumor cells are shown in green, macrophages in red, and blood vessels in yellow. Scale bar = 50 μm, time is indicated in hours:minutes, T_i_ = 20 min, frame rate = 4 fps.

